# Bracket Coding: An Emergent Balance Between Temporal Integration and Segregation in Early Visual Population Activity

**DOI:** 10.64898/2026.05.31.729124

**Authors:** Toktam Samiei, Hafiz Fareed Ahmed, Edward Zagha, Erfan Nozari

## Abstract

Despite over a century of research into the neural code, the fundamental principles by which the brain encodes sensory information remain debated. In this study we provide converging evidence for the presence of a dynamic, fast-switching integration of rate and temporal coding in the thalamus, primary visual cortex, and higher-order visual cortical areas of mice viewing an array of visual stimuli. This scheme is primarily characterized by the presence of distinct, temporally coordinated “bracket”s that tile the duration of each trial, are rate-coded within, and are separated by boundaries that are precisely-timed and synchronized across the population. Using large-scale Neuropixels recordings from the Allen Institute Visual Coding dataset, we provide evidence for the robustness and generality of bracket coding across several visual tasks and brain regions, as well as its optimality for information decoding, functional relevance for information representation, pronounced hierarchical organization, long-range bottom-up synchrony across visual regions, and coherence with low-frequency local field oscillations. These findings were all subsequently validated in a second, independent dataset provided by the International Brain Laboratory consortium. Finally, we provide a computational model that can serve as a potential mechanism for the generation of bracket-coded population spiking activity. Together, our results demonstrate the presence of a novel form of sensory information encoding in the brain, with broad implications for neuroscience and neuroengineering.

## Introduction

Understanding how the brain encodes information in patterns of neural activity is one of the central questions of neuroscience. Across sensory, motor, and cognitive systems, neurons transform external inputs and internal states into sequences of spikes that must ultimately support perception, decision-making, and behavior. Yet despite decades of progress, the principles by which these spikes represent information remain debated. In particular, which aspects of spiking activity are meaningful to downstream circuits, and over what timescales is that meaning organized, is only partially understood. Resolving this question is essential not only for understanding biological computation, but also for interpreting large-scale neural recordings, constructing accurate decoding models, and building biologically grounded brain–computer interfaces and artificial intelligence systems [1, 5, 7, 11, 29, 35].

Historically, the notion of rate coding—that information is carried only by the number of spikes emitted over a temporal window—has remained influential, owing mainly to its simplicity and robustness for information decoding [1, 11, 35, 37, 38]. Early electrophysiological work established a close relationship between stimulus intensity and impulse frequency in sensory afferents [1], and later studies showed that average firing rates could explain major aspects of stimulus selectivity and behavioral readout across many systems [18, 20, 26]. More broadly, rate-based population coding has offered an appealing framework in which downstream neurons can recover stable representations by integrating activity over time [4, 29, 32, 33].

At the same time, a large body of work has shown that in many systems, the precise timing of spikes, as well as temporal relations across neurons, carries information beyond that available from rates alone [8,35,43,45]. This includes the early demonstration of millisecond-scale reliability in cortical spike timing under fluctuating inputs [25], as well as various studies showing that short spike sequences and first-spike latencies can carry substantial information about sensory stimuli [6, 12, 13, 19, 30, 34, 44]. Beyond sensory cortex, related forms of temporal coding have been observed in prefrontal short-term memory, where object information is organized by oscillatory phase [17,22,39], and in the hippocampus, where theta phase precession provides a temporal code for position within place fields [9, 14, 27, 41]. Such observations have motivated a broader view in which temporal precision supports rapid discrimination, dynamic sensory processing, and efficient representation of behaviorally relevant events [8, 28, 43].

This contrast between rate-based and time-based accounts has motivated a growing set of hybrid frameworks in which neural coding is understood as an interaction between firing rates and temporal organization rather than a strict choice between the two. Reviews of sensory and population coding increasingly emphasize that meaningful representations may emerge jointly from rates, correlations, spike timing, oscillatory phase, and structured dynamics across multiple temporal scales [4, 16, 21, 22, 28, 29, 31]. One influential example is packet coding, which proposes that cortical activity is organized into transient packets of population activity within which both rate and fine temporal structure contribute to representation [23, 24]. More generally, such perspectives suggest that stable information may persist over limited temporal intervals, while transitions between representational states are marked by temporally coordinated changes in activity. In this sense, rate and temporal coding may not be competing theories, but complementary components of a richer dynamical code.

Despite this progress, the field still lacks a clear account of how rate-based stability and temporally precise coordination are integrated within ongoing population activity. Addressing this gap not only requires a better understanding of what variables carry information, but also how that information is temporally organized. In this work, we address this question by demonstrating converging evidence in support of a new coding scheme, called bracket coding. The hallmark of bracket coding is a balanced coexistence of rate and temporal coding, where visual population activity is organized into temporally bounded intervals within which information remains relatively stable, while transitions between intervals are marked by precise coordinated temporal structure. Bracket coding therefore provides a principled way to reconcile rate-based and temporal accounts of neural coding in early visual processing, and offers a general perspective on how neural populations may combine stability and precision to support fast and reliable computation.

## Results

### Spike brackets as intervals of optimal integration in population spiking activity

To better understand the *dynamic structure* of information encoding in population spiking activity, we developed a framework consisting of two components: (1) an end-to-end decoding block that measures the amount of information present in a data stream (population spiking activity) by how accurately a normative decoder can decode the identity of a visual stimulus from that data stream, and (2) a sweeping-integration block right *before* the decoder that performs local temporal integration over increasingly larger windows (Figure 1a). Motivated by our recent theoretical analyses [2], our approach is based on the premise that when integration over a window improves decoding accuracy, population spikes within that window encode similar information (shared signal *>* shared noise). In contrast, when integrating spikes over an interval degrades accuracy, population spikes within that interval encode *dissimilar* information (shared signal *<* shared noise).

**Figure 1.**
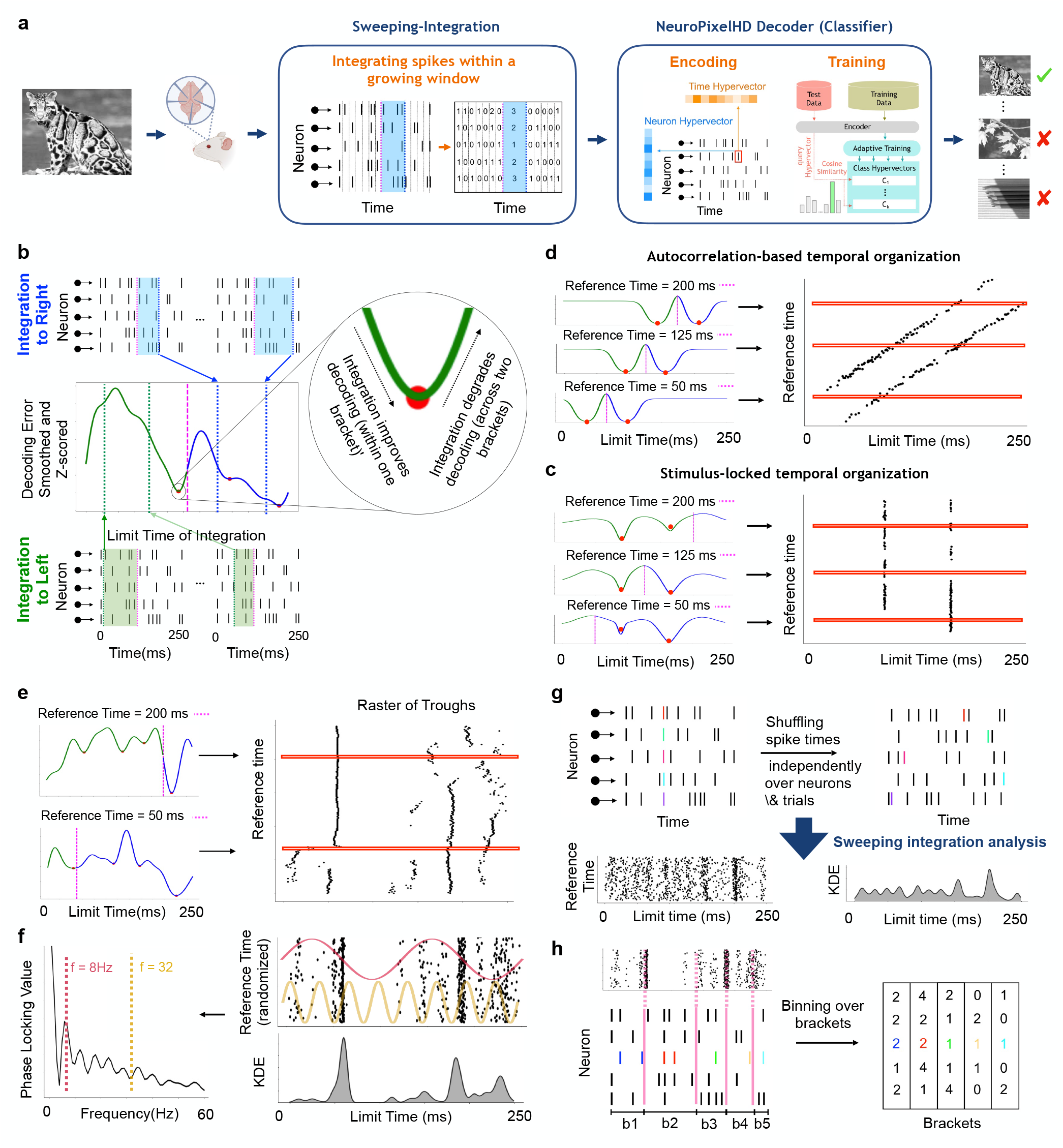
Overview of bracket coding in populations of spiking neurons. **(a)** The computational framework for identifying intervals of optimal integration in population spiking activity. Single-unit population spikes (Allen Institute Visual Coding - Neuropixels data [40], see Methods) were first passed through a sweeping integration block before being fed to a normative decoder. The latter has been developed and described in prior work [36]. The former integrated (‘binned’) spikes within a growing window that spanned between a fixed ‘reference time’ (RT, magenta) and a moving ‘limit time’ (LT, blue). This integration can both decrease or increase decoding error, depending on whether the integrated spikes carried statistically similar to dissimilar information about the stimulus. **(b)** For each fixed RT, sweeping integration was performed in both directions, towards left (top panel, blue) and right (bottom panel, green). We then tracked the trajectory of the resulting decoding error, as a function of the LT of integration. The troughs (local minima) of this trajectory, by definition, partition the trial duration into intervals where integration within one interval monotonically improves decoding but integration over distinct intervals starts to increase the error (inset). **(c**,**d)** Cartoon schematic of two alternative possibilities for the temporal organization of error troughs. The error trajectories on the left side of each panel result from repeating the process explained in (b) for each RT between trial start (0ms) and trial end (250ms). The result is then summarized via the trough rasters on the right, where each dot represents one trough and each row corresponds to one error trajectory (i.e., one RT). If the similarity of information represented at two time points depends primarily on their temporal distance (autocorrelation-based organization, panel (c)), error troughs should shift together with the reference time (left), resulting in diagonal bands in the trough raster (right). If, however, similarity of information represented at two time points depends primarily on their absolute timing relative to stimulus onset (stimulus-locked organization, panel (d)), error troughs should occur at fixed limit times regardless of the reference time of integration (left), giving rise to vertical bands in the trough raster (right). **(e)** Example error curves and corresponding trough raster obtained from real data. Trough locations are largely organized vertically, rather than diagonally. **(f)** To quantify the strength of vertical alignment, each raster of troughs was summarized into a density over time, visualized as a kernel density estimate (KDE, bottom left) as well as a strip plot (top left, equivalent to the raster in (e) shuffled over reference times). This density exhibits pronounced and isolated peaks, hereafter referred to as *bracket boundaries*, which reflect a remarkable consistency in the location of error troughs across different (even far-apart) reference times. To quantify this consistency, we computed the phase locking value (PLV) between the error troughs and sinusoids of different frequencies between 0-60Hz (right). **(g)** To statistically test the significance of PLVs (as well as other statistics), we generated surrogate data by time-shuffling the spike times of all neurons. This process destroys all dynamic structures in information encoding while preserving static rates. Surrogate data was passed through the same analysis pipeline as real data, and their resulting statistics were compared at the population level. **(h)** Schematic showing final discrete bracket boundaries obtained as the prominent peaks of the trough density estimated in (f). These brackets can then be used for an optimal (non-uniform) binning of population spikes as shown.

Central to this process, the sweeping-integration block starts from raw data represented at full (1 ms) resolution, and locally integrates each neuron’s spikes over a growing *integration window* (Figure 1b). This window spans the interval between a fixed time and a moving time (relative to stimulus onset), to which we refer as the *reference time* (RT) and *limit time* (LT), respectively. As we gradually expand the integration window, we record the resulting decoding error, and the process is repeated for all RTs throughout the trial period. Depending on the relative position of the RT and LT, spikes are integrated both from preceding time points (integration to the left, LT *<* RT) and, separately, from subsequent times (integration to the right, LT *>* RT). By performing both procedures for each RT, we obtain an *error trajectory* that reflects the local similarity of information around each RT in the population spiking activity.

We then computed the times of switching in information content via the *troughs (local minima)* of each error trajectory (Figure 1b). Troughs of an error curve, by definition, mark the transition points between an interval where integration has been lowering decoding error, and the subsequent interval whose additional integration starts to increase the error. In other words, each trough marks a point in time where similar spikes have been integrated, and extending the window further starts to increase error by mixing-in dissimilar spikes from the subsequent interval. As such, the troughs of each error curve *partition* the trial duration into intervals of locally-similar information. This partitioning is, however, relative to the RT from which integration started. If similarity in the information content of two time points varies only as a function of their distance (i.e., the autocorrelation function), troughs should move together with the RT (Figure 1c). If, on the other hand, similarity in information content varies based on fixed intervals (time-locked to stimulus onset), troughs would remain consistent regardless of the RT (Figure 1d). To distinguish between these possibilities, we computed error trajectories for every RT across the trial, and summarized their troughs in the form of a *trough raster* across all RTs and trough locations.

As shown in Figure 1e for a sample subject, we observed the location of error troughs to be highly consistent (time-locked to stimulus onset) and independent of the reference time from which integration started. This stability provides strong evidence for coordinated population-level timing during information encoding. To quantify the statistical significance of this temporal alignment, we computed the phase-locking value (PLV) between trough locations and sinusoidal signals across different frequencies (Figure 1f). PLV measures the presence of periodic temporal alignment in the distribution of troughs, with strongly-aligned structures producing higher PLVs and uniformly scattered troughs yielding PLV≃ 0.

To obtain a surrogate null distribution, we then repeated the same procedure on temporally-shuffled spike trains whereby spike times were randomly shuffled over time, independently across neurons and trials (Figure 1g). Compared to real data, the shuffled controls produced boundaries that lacked strong temporal consistency across trials. This difference is evident in the kernel density estimates (KDEs) of the distributions of trough locations, which is relatively flat in shuffled but clustered around specific time points with pronounced peaks in real data, as well as the distributions of real and shuffled PLVs (Figure 2).

**Figure 2.**
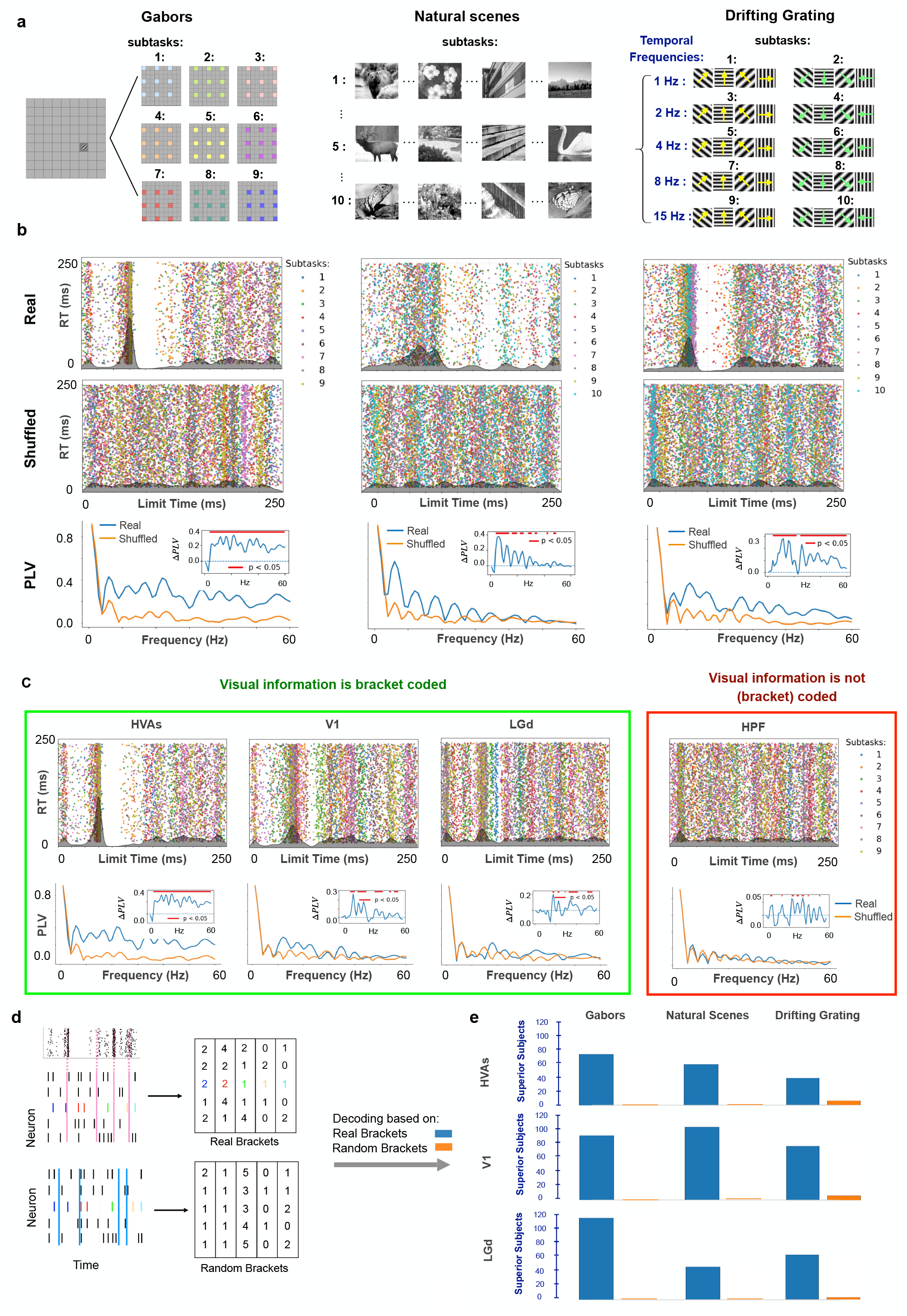
Generality of bracket coding across brain regions and visual tasks. **(a)** Neuropixels data from three visual tasks in the Allen Institute Visual Coding–Neuropixels dataset [40] were analyzed in this study: Gabor patches, natural scenes, and drifting gratings. Each task was divided into multiple classification problems (subtasks) by grouping trials into disjoint sets of classes based on stimulus condition. In the localization of Gabor patches, we divided the 81 possible positions of the stimulus into 9 sets (subtasks) of 9 locations each uniformly distributed across the visual field. In the identification of natural scenes, the 118 image classes were partitioned into ten sets of ~ 12 images each. In drifting gratings, we divided the 8 possible grating orientations into two sets, and further separated them based on their temporal frequencies into 10 total subtasks. For each subtask, the sweeping-integration analysis (Figure 1) was applied independently and the resulting trough rasters were combined (overlaid) across subtasks. **(b)** Example trough rasters (colored dots), trough densities (gray), and PLVs of real and shuffled data in higher visual areas (HVAs) across the three tasks. As noted, the trough rasters are overlaid across all subtasks within each task. Each column shows data for one subject (cf. Figure 4 for group-level statistics). Insets show the Δ*PLV* = *PLV*_real_ − *PLV*_shuffled_, with red segments indicating statistically significant differences that demonstrate the presence of bracket coding (*p <* 0.05, one-sided bootstrap test, FDR corrected for multiple comparisons). **(c)** Presence of bracket coding across brain regions. Example raster plots and PLV spectra are shown for HVAs, primary visual cortex (V1), dorsal lateral geniculate complex (LGd), and the hippocampal formation (HPF) during the viewing of Gabor patches. The latter serves as a control region outside the visual pathway. Details parallel those in panel (b). Significant evidence of bracket coding is observed across all visual regions (HVAs, V1, and LGd) but not in HPF. **(d)** Schematic comparison between decoding based on the real bracket boundaries extracted from population activity (top) and that based on randomly selected boundaries (bottom). In both cases, integrated spike trains within the respective time intervals were used for decoding. **(e)** Summary across tasks and regions of the number of subject-subtask combinations in which decoding based on each scheme (real brackets or random boundaries) statistically significantly outperformed decoding based on the other. Across all tasks and regions, decoding based on real brackets significantly outperformed decoding based on random boundaries.

Thus, in conclusion, the observed temporal alignment in the trough rasters reveals fixed temporal boundaries that partition neural activity into intervals, referred to as *‘brackets’* hereafter. Accordingly, we use *‘bracket coding’* to refer to the hybrid coding scheme characterized primarily by the partitioning of the duration of each trial into intervals (brackets) such that (1) within each interval spikes encode highly similar information (rate coding), while (2) the *boundaries* between intervals remain stable from trial to trial and time-locked to stimulus onset (temporal coding). In the remainder of this article we test the robustness and generality of bracket coding, as well as characterize its properties and functional relevance.

### Bracket coding is observed across brain regions and visual tasks

We performed the sweeping integration analysis (Figure 1) on data from n = 10 mice across three visual tasks—passive viewing of Gabor patches, natural scenes, and drifting gratings—in the Allen Institute Visual Coding– Neuropixels dataset. We further partitioned the available trials for each subject and task into multiple disjoint *subtasks* to increase statistical power and better distinguish true brackets from bracket-like structures that may arise in shuffled data due to finite samples (Figure 2a). Each subtask corresponds to a separate classification problem, for which a decoder is trained independently for each subject. This was done based on the premise that, if the observed brackets reflect true temporal structure in the underlying population activity, they must remain consistent across independent sets of trials (subtasks). If, in contrast, the observed brackets reflect random fluctuations in finite-sample statistics, they should occur at independent times across independent subtasks. We thus overlaid trough rasters across all subtasks, and computed a single trough density and the corresponding PLV for each subject. This PLV was then compared against its null distribution obtained from surrogate shuffled data and the comparison was used to test the presence of bracket coding in each region and task.

Across all subjects and tasks, we observed significant evidence that visual information is bracket-coded throughout different stages of visual processing (Figure 2b,c). As shown in Figure 2b for higher visual areas (HVAs, cf. Methods), PLVs computed from real data (blue) were significantly higher than those obtained from shuffled data (orange) (*p <* 0.05 at most frequencies, one-sided bootstrap test, FDR corrected for multiple comparisons). This difference was observed across all three visual tasks, reinforcing the presence of robust temporal alignment of bracket boundaries in real data that is absent in the shuffled controls.

We next examined whether evidence of bracket coding is present across different brain regions. As seen from Figure 2c, areas directly involved in visual processing—primary visual cortex (V1), HVAs, and the dorsal lateral geniculate complex of the thalamus (LGd)—showed clear evidence of bracket coding (*p <* 0.05 at most frequencies, one-sided bootstrap test, FDR corrected for multiple comparisons). In contrast, PLVs computed by applying the same procedure to data from the hippocampal formation (HPF) exhibited no significant difference between real and shuffled data, confirming the absence of bracket coding in HPF, where low-level visual information pursued herein is not expected to be encoded.

### Optimality of bracket-based decoding for information retrieval

Given the robustness of our findings across tasks, regions, and datasets, we hypothesize that bracket coding may reflect an optimal coding regime for spike-based communication. To test this, we compared decoding of visual information based on real versus randomly generated brackets (Figure 2d). Indeed, when neural activity was integrated within the empirically identified brackets and segregated across successive brackets, decoding performance was higher than when the same was performed using random boundaries, consistently across brain regions and visual tasks (Figure 2e). This result highlights that the advantage of bracket-based decoding does not arise from access to more data, but from integrating activity according to boundaries that reflect genuine structure within population spiking activity.

### Distinct dimensions of sensory information are encoded within different brackets

Having established the presence of bracket coding as a statistical regularity in the dynamics of normative decoding error, we next sought to examine its functional relevance by analyzing the information content of population spiking activity across brackets. To this end, we performed a temporal dissimilarity analysis by considering two adjacent, non-overlapping windows and evaluating whether they carry functionally distinct information. We slid the two windows across the entire trial and quantified the dissimilarity between their decoding accuracy profiles using the Euclidean distance between their respective vectors of per-class decoding accuracy. The resulting dissimilarity curve was then compared against the kernel density estimate (KDE) of the bracket boundary distributions of each subject (Figure 3a). We hypothesize that, if bracket boundaries mark transitions between distinct informational states, peaks in the dissimilarity curve should align with bracket boundaries, whereas the two should remain statistically independent otherwise.

**Figure 3.**
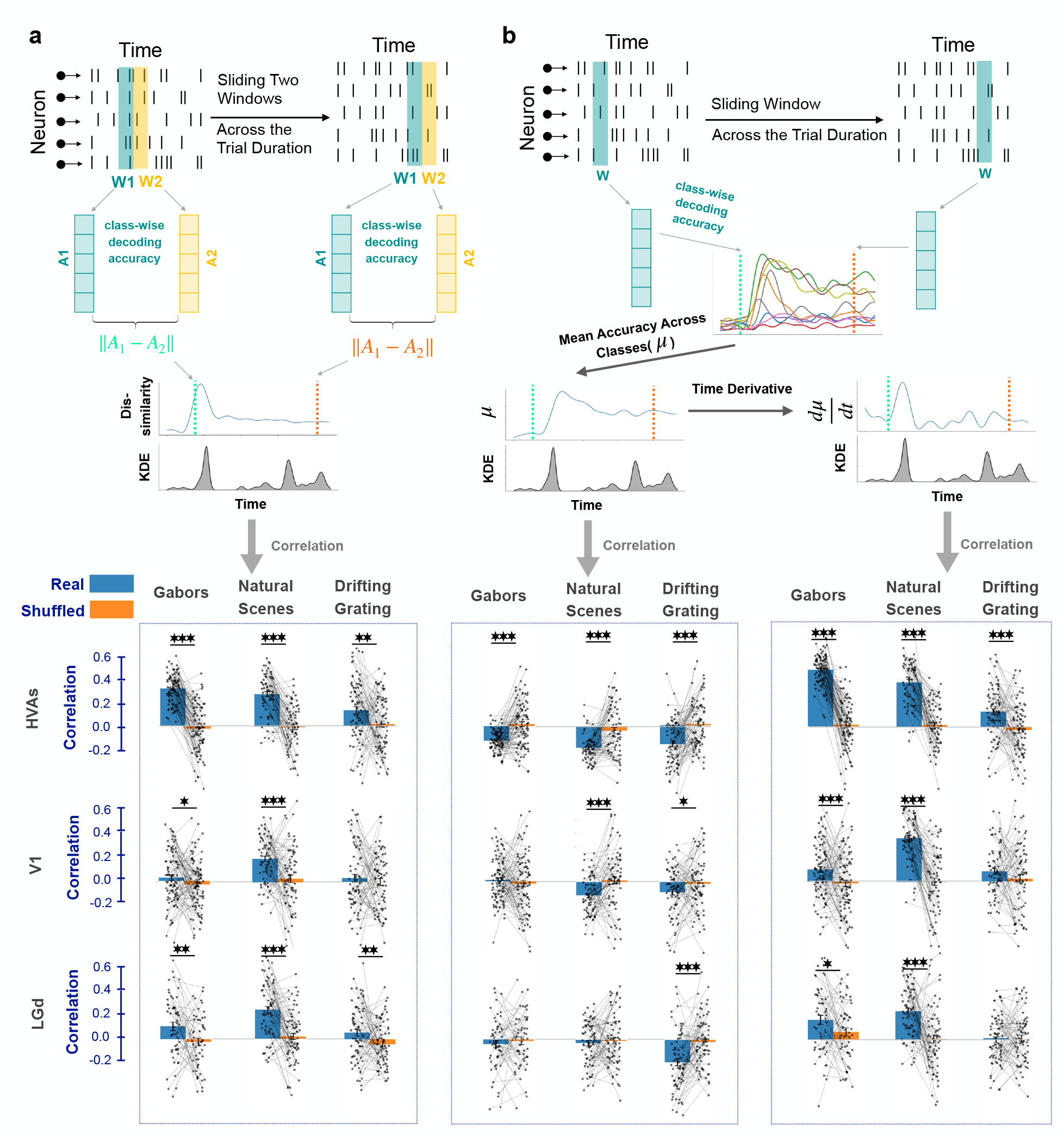
Distinct dimensions of sensory information are encoded within different brackets. **(a)** Temporal dissimilarity analysis. Two adjacent, non-overlapping windows (W1 and W2, 15ms-wide each) were slid across the trial duration. For each window, population activity was used to decode stimulus identity, producing class-wise decoding accuracy vectors (*A*_1_ and *A*_2_, respectively). The dissimilarity between the latter ∥*A*_1_ − *A*_2_∥ was then calculated at each sliding step and the resulting time-series was correlated against the kernel density estimate (KDE) of bracket boundaries. The same procedure was repeated on shuffled data. Bottom panel shows the distribution of the resulting correlation coefficients in real and shuffled data across all tasks and brain regions, with the strongest effects in higher visual areas (HVA) and during the viewing of natural scenes. Each dot in the strip plots represents one subtask from one subject. **(b)** Temporal multiplexing analysis. A single 15-ms window (W) was slid throughout the trial duration and the population spiking activity within this window was used to decode the presented stimulus. The time series of mean accuracy was then correlated against the density of bracket boundaries. The same was done for the time derivative of mean accuracy, and both analyses were repeated on shuffled data. Across the majority of tasks and regions, mean accuracy and its derivative showed significant negative and positive correlations with bracket boundaries, respectively, indicating rapid reorganization of decoding structure at bracket boundaries. All correlations were near zero on shuffled data. Similar to (a), effects are strongest in HVA and during the viewing of natural scenes.

Across nearly all tasks and brain regions, we observed significant positive correlations between bracket boundaries and the dissimilarity of information carried by adjacent windows (Figure 3a). The effect was particularly pronounced in HVAs, which exhibited the strongest and most consistent correlations across all tasks. Among the task conditions, natural scene viewing showed the strongest correlations between bracket boundaries and information dissimilarity, indicating the presence of particularly strong transitions in decodable information across bracket boundaries in this task. Importantly, the same analysis performed on shuffled data resulted in near-zero correlations in all cases which, in nearly all tasks and brain regions, were statistically smaller than correlations in real data (Wilcoxon signed-rank test,*p <* 0.05). Together, these results indicate that bracket boundaries reliably coincide with abrupt transitions in the informational content of population activity, supporting the view that adjacent brackets encode functionally distinct neuronal representations.

### Temporal multiplexing underlies the emergence of bracket structure

Taking this analysis further, we next sought to characterize the exact differences that exist in information content across brackets. We hypothesized that, if bracket boundaries correspond to transitions between distinct informational states, the accuracy by which different stimulus classes can be decoded should systematically reorganize around those boundaries. In other words, population activity should *not* encode all stimulus classes uniformly across the trial, but rather in a way that each stimulus class is associated with a distinct temporal information profile (Supplementary Fig. S1). We refer to this phenomenon as *temporal multiplexing* in an operational sense: encoded stimulus information is distributed across time in a class dependent manner, so that the pattern of decodability across classes changes over the course of the trial. This usage is motivated by prior work on multiplexed sensory codes across temporal scales [28], but here refers specifically to time-varying, class-wise decoding structure. Under this interpretation, bracket boundaries should correspond to moments when the population representation reorganizes, which may appear as transient changes in the average decodability across classes and/or rapid changes in that average over time.

To examine this, we computed the class-wise decoding accuracy within a small 15ms-sliding window throughout the trial duration. This yielded, at each time point (for each window), a vector of per-class decoding accuracies from which we derived two summary time-series: (1) the mean decoding accuracy across classes, and (2) the temporal derivative of this mean. We then compared the temporal dynamics of these time-series against the dynamics of bracket boundaries (Figure 3b).

The results revealed significant correlations between decoding statistics and bracket boundaries across tasks and brain regions (Figure 3b). Most notably, the temporal derivative of the mean decoding accuracy showed a significant *positive correlation* with bracket boundaries in most tasks and brain regions, indicating that bracket boundaries reliably coincide with moments of rapid change in information content (Wilcoxon signed-rank test, *p <* 0.05). In contrast, a significant *negative correlation* exists between the mean decoding accuracy itself and the bracket boundaries in most tasks and regions, indicating that average decoding performance tends to drop near bracket boundaries (Wilcoxon signed-rank test, *p <* 0.05). Consistent with the temporal dissimilarity analysis, the strongest and most statistically significant multiplexing effects (both mean and derivative of the mean) were observed in the natural scenes task (across all areas) and in HVAs (across all tasks). Together, these patterns imply the presence of rapid, transient disruptions of one informational regime before the emergence of the next one separated by bracket boundaries. In other words, decoding accuracy briefly drops while the representational structure shifts, after which a new relatively stable informational regime emerges within the subsequent bracket.

### Bracket coding follows established hierarchical gradients along the stages of thalamocortical visual processing

Given the functional relevance of bracket coding for visual information processing, we hypothesized that its emergence should vary systematically across brain regions in accordance with the anatomical and functional hierarchy of the visual system. To test this, we first examined the strength of bracket coding measured by the largest difference between the PLVs computed from real data and shuffled surrogates (max_*f*_ Δ*PLV* (*f*)), as well as the number of frequencies (*N*_*sig,freq*_) at which this difference is significant (*p <* 0.05, paired *t*-test, FDR-corrected for multiple comparisons)(Figure 4a). Consistent with our hypothesis, we observed a clear hierarchical trend across all subjects and tasks, with both metrics increasing along the visual hierarchy from LGd to V1 to HVAs. In contrast, HPF (control region) showed little to no significant PLVs in any of the tasks, confirming the specificity of this hierarchical organization to visual regions.

**Figure 4.**
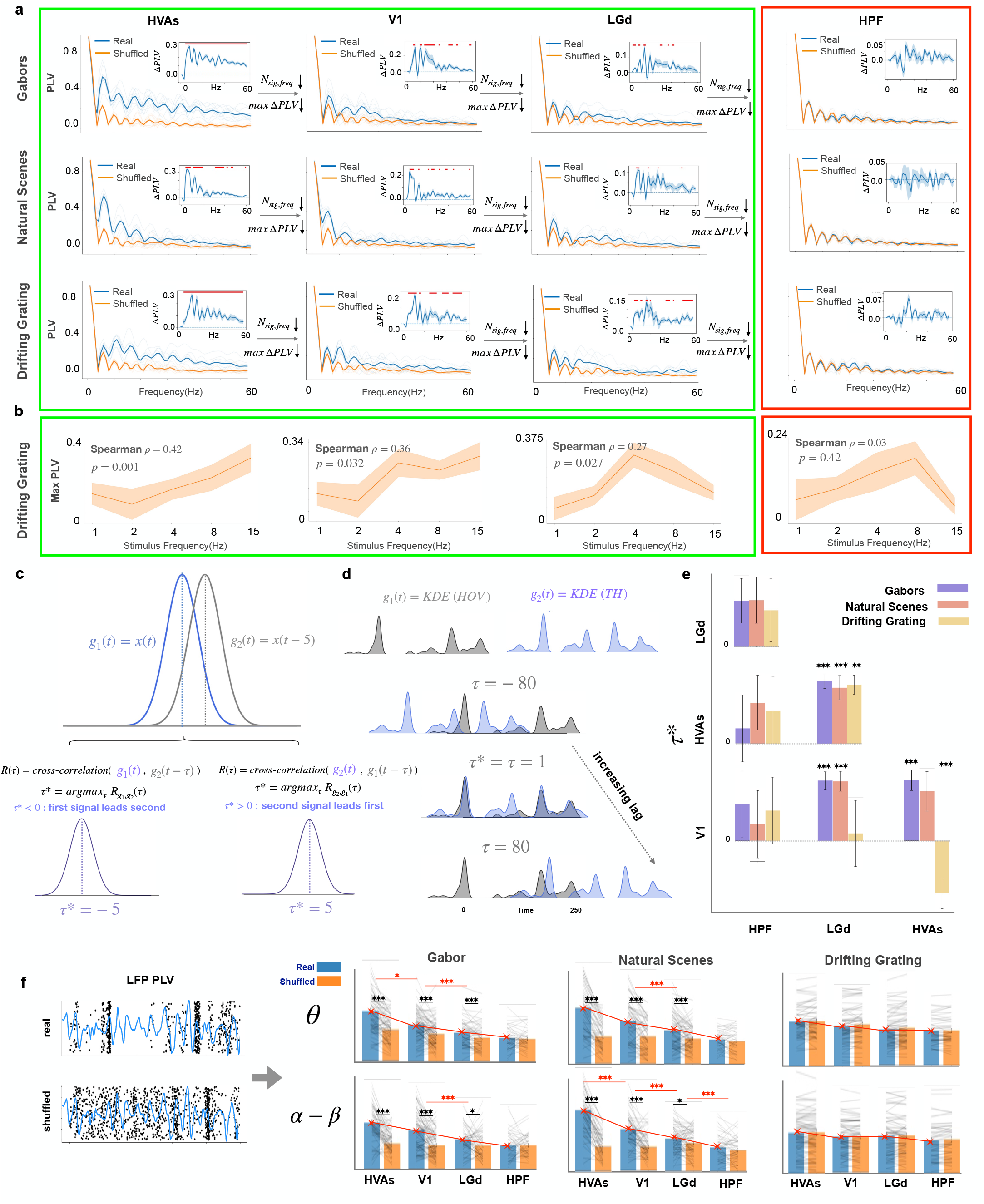
Hierarchical organization of bracket coding along the thalamocortical visual system. **(a)** Average phase-locking value (PLV) spectra of bracket boundaries in real data and shuffled controls across all tasks (Gabor, Natural Scenes, and Drifting Gratings) and brain regions (HVA, V1, LGd, and HPF). For each condition, PLV was computed at each frequency and averaged across sessions. Insets summarize two quantitative metrics extracted from each spectrum: the number of significant frequencies (*N*_sig,freq_), defined as frequencies at which real PLV significantly exceeds shuffled PLV (FDR-corrected *p <* 0.05, paired t-test), and the maximum PLV difference (max ΔPLV), defined as the maximum value of the difference between average real and shuffled PLVs across all frequencies. Across tasks, both metrics reveal a consistent hierarchy in which bracket coding gradually strengthens as we go up the visual hierarchy, and specificity to visual regions (not present in HPF). **(b)** Relationship between stimulus temporal frequency and maximum Δ*PLV* during the viewing of drifting gratings. For each drifting frequency in the drifting grating task, we first computed ΔPLV = PLV_real_ − PLV_shuffled_ across all analyzed PLV frequencies, then took the maximum ΔPLV across those frequencies, and finally correlated this maximum value with the drifting frequency. Significant and hierarchically organized correlations between the two exist in visual regions but not in HPF. **(c)** Schematic illustration of the cross-correlation procedure used to estimate temporal lags between pairs of signals. Two example signals with a temporal offset produce a cross-correlation function whose peak defines the optimal lag *τ*\*. Positive and negative values of *τ*\* indicate which signal leads in time. **(d)** Schematic illustration of applying the cross-correlation analysis in (c) to kernel density estimates (KDEs) of bracket boundaries across brain regions. Temporal shifts between KDE signals were computed to estimate relative timing of bracket structures across regions. **(e)** Summary of cross-regional temporal relationships inferred from the cross-correlation analysis in (d) across tasks and brain regions. The regions depicted on the vertical and horizontal axes were treated as the first and second signals, respectively. Therefore, following the sign convention in (c), positive *τ*\* values indicate that the second region leads the first. The estimated lags reveal a hierarchical organization in which LGd brackets significantly lead those in V1, which in turn significantly lead those in HVA (Paired Wilcoxon signed-rank test on cross-correlation values across pairs (*p <* 0.05)). This is consistent with a feedforward propagation of bracket-coded structure along the visual hierarchy. No significant lead-lag relationships exist between HPF and other regions. **(f)** The analysis of phase-locking between bracket boundaries (represented by raster of troughs, black) and narrow-band local field potentials (LFPs, blue). PLVs were computed between the two in *θ* and *α*–*β* bands, both in real and shuffled data. Results demonstrate significant bracket–LFP coupling across tasks, with increasing strength along the visual hierarchy (HVA *>* V1 *>* LGd). Across all panels\**p <* 0.05, ^*\*\*\**^*p <* 0.001 and error bars show 1 s.e.m.

During the viewing of drifting gratings, the strength of bracket coding is further modulated, with a hierarchical gradient, by the temporal drifting frequency of the stimulus. As seen in Figure 4b, max_*f*_ Δ*PLV* (*f*) (now computed separately for each drifting frequency) is significantly modulated by the stimulus frequency across all visual regions (*p <* 0.05, one-sided permutation test on Spearman’s *ρ*) This shows, in particular, that the temporal structure of the stimulus influences the rhythmic consistency of bracket boundaries. Further, this modulation follows a hierarchical ordering, where the correlation is strongest in HVAs, followed by V1 and LGd, and insignificant in HPF. This hierarchy is particularly notable because the influence of stimulus timing *increases* along the visual hierarchy, and thus minimizes the possibility that bracket boundaries merely reflect the timing of stimulus presentation. Instead, while bracket timing is modulated by the temporal structure of the stimulus, it is an emergent property of network dynamics that becomes progressively amplified and refined along the visual processing hierarchy.

### Bracket boundaries are long range-synchronized across stages of visual processing

To determine whether bracket timing is coordinated across brain regions in a manner consistent with hierarchical information flow, we examined lead–lag relationships between bracket boundaries across the visual pathway. To this end, we cross-correlated bracket boundary KDEs between each pair of brain regions, quantifying their temporal alignment across different amounts of relative time lag (Figure 4c). The lag at which the cross-correlation reached its maximum was then extracted and used to test the presence of consistent lead–lag relationships among regions at the group level (Figure 4d). Across all the considered visual tasks, we observed significant, predominantly bottom-up long-range synchrony of bracket boundaries among thalamocortical visual regions (Figure 4e). Out of 9 total pairwise comparisons, we observed statistically significant long-range synchrony between 7 pairs (*p <* 0.01, Wilcoxon signed-rank test), with 6 following a bottom-up temporal order (LGd leading V1 leading HVAs). These lead–lag relationships were specific to visual regions, with no significant long-range synchrony observed between the HPF other regions in either task. This specificity further supports the functional relevance of the observed long-range relationships among visual regions. Together, these findings support the hypothesis that bracket boundaries participate in long-range temporal coordination of neural population activity, and is consistent with hierarchical information flow along the thalamocortical pathway.

### Bracket boundaries are phase-locked to low-frequency local field potential oscillations

We next tested the presence of temporal coherence between the timing of bracket boundaries and the local field potential (LFP) in each region, which could serve as a potential mechanism for coordinating spiking activity across large populations of neurons. Similar to PLVs described so far for phase-locking to single-frequency oscillations (Figure 1d),we computed the PLV between bracket boundaries and narrow-band LFP oscillations by band-pass filtering each region’s mean LFP in the theta (4–8 Hz) and alpha–beta (8–30 Hz) bands and computing the PLVs of real and shuffled error troughs to each narrow-band LFP. Across all visual regions and both frequency bands, we observed significantly stronger phase locking of bracket boundaries to LFPs in real data compared to shuffled surrogates during the viewing of Gabors and Natural Scenes. Further, the strength of this phase locking exhibited a clear hierarchical gradient along the visual pathway in all tasks: PLV values increased from LGd to V1 and reached their highest levels in HVAs. This progressive increase suggests that the formation of bracket coding becomes stronger as visual information propagates through successive stages of the thalamocortical visual hierarchy, consistent with the proposition that bracket timing emerges from the intrinsic dynamics of cortical population activity rather than raw sensory input. Consistent with prior findings, HPF showed no significant phase locking to LFP in either band or task. Thus, in summary, we observe a significant and consistent hierarchical organization in several independent aspects of bracket coding, supporting its emergence from local network dynamics and its functional relevance as a means of inter-regional information transfer.

### Weaker but significant bracket coding of visual information during sensorimotor discrimination

To test the generality of bracket coding beyond the Allen Institute passive visual tasks, we repeated the same analysis pipeline to Neuropixels data from *n* = 114 mice during visual decision making, available as part of the International Brain Laboratory (IBL) project [3]. In this task, head-fixed mice were presented with a Gabor patch of fixed orientation and varying contrast on either side of the screen, and rotated a wheel to center the stimulus within a 60-s response window to obtain reward (Figure 5a, see [3] for details). We used trials with the highest levels of contrast (100% and 25%) and partitioned them into two subtasks for end-to-end decoding (Figure 5a). Given the large cohort available in the IBL dataset but the significantly lower number of ‘good’ neurons available per mouse, we pooled neuronal activity across randomly-selected groups of *k* = 1, 3, 5, 10, or 15 mice into ‘hypermice’ whose neurons were then treated as if simultaneously recorded (Figure 5b and Methods). Notably, although the emergence of bracket coding in the resulting hypermice would require a universal (shared) timing of brackets across the constituent mice, we observed an increase in the strength of bracket coding (max_*f*_ Δ*PLV* (*f*)) with increased *k* (Figure 5c, see also Discussion). Based on this analysis we selected *k* = 15, resulting in *n* = 15 hypermice that were used in the subsequent analyses.

**Figure 5.**
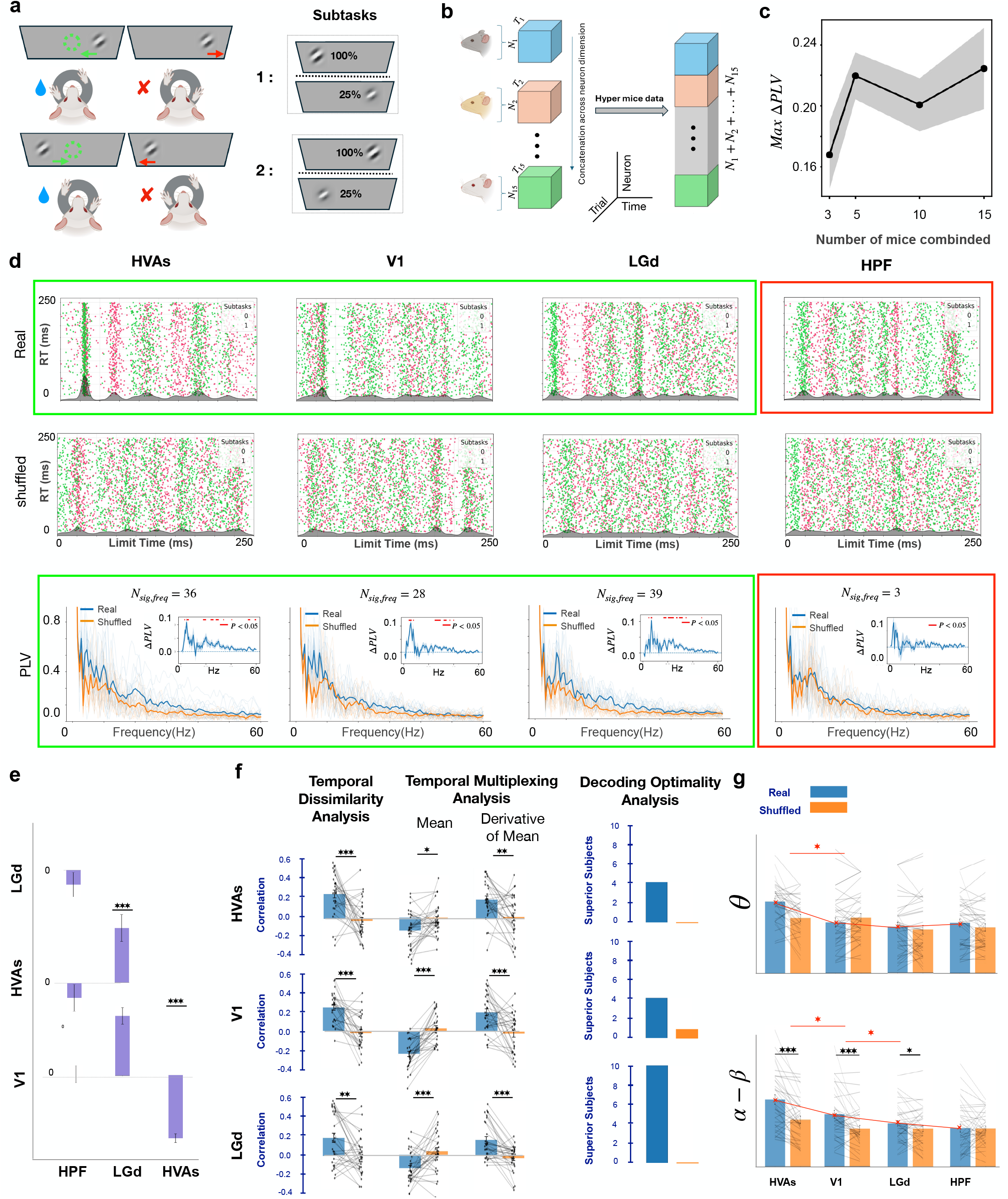
Bracket coding of visual information during visual decision making. (**a)** Schematic of the IBL visual decision-making task. Head-fixed mice were presented with a Gabor stimulus at varying contrast levels and reported its location via wheel movement. Construction of classification problems (subtasks) using trials from high-contrast conditions (100% and 25%) is shown on the right. **(b)** Neurons from groups of *k* = 15 randomly-selected mice were pooled into ‘hypermice’ (treated as simultaneously recorded). The number of trials for each hypermouse was limited to the minimum available across all constituent mice (*min* {*T*_1_, *T*_2_, …, *T*_15_}). **(c)** Strength of bracket coding (max ΔPLV) as a function of the number of pooled mice sessions (*k*). The former initially increases with *k* and then saturates. **(d)** Example rasters of error troughs (colored dots, color-coded for subtask) and their KDEs (gray) for real and temporally shuffled data across the same brain regions as in Allen data (HVAs, V1, LGd, and HPF). Corresponding real and shuffled single-frequency PLV spectra are shown in the bottom row for all subjects (hypermice). Insets show Δ*PLV* = *PLV*_real_ − *PLV*_shuffled_ and number *N*_*sig,freq*_ of frequencies at which this difference is significant (*p <* 0.05, one-sided Wilcoxon signed-rank test, FDR-corrected for multiple comparisons). **(e)** Results of cross-regional long-range synchrony analysis. Details parallel those in Figure 4c-e. While the lead-lag order in two of the three visual region pairs is on average consistent with the bottom-up direction observed in Allen Institute data, only the one between LGd and HVAs is significant (Paired Wilcoxon signed-rank test on cross-correlation values across pairs (*p <* 0.05)), and the temporal order between V1 and HVAs is significant in the reverse (top-down) direction. HPF still shows no significant lead-lag relationships. **(f)** Summary across tasks and regions of temporal dissimilarity, temporal multiplexing, and decoding optimality analyses. Analysis details parallel those in Figures 3 and 2e. Across tasks and regions, temporal dissimilarity and temporal multiplexing correlations were statistically significant and consistent in sign with the Allen Institute results. Decoding based on real brackets also significantly outperformed decoding based on random boundaries, matching the pattern observed in the Allen Institute data. **(g)** Phase-locking between bracket boundaries and local field potentials (LFP). Details parallel those in Figure 4f. We still observe significant bracket-field coherence with a hierarchical gradient, particularly in the *α*–*β* band. Across all panels\**p <* 0.05, ^*\*\**^*p <* 0.01, ^*\*\*\**^*p <* 0.001 and error bars show 1 s.e.m.

As seen from Figure 5d, we observe evidence of bracket coding, namely, significantly larger PLVs of bracket boundaries at several frequencies in all subjects in real data compared to shuffled surrogates, during sensorimotor decision making as well. The strength of bracket coding is notably weaker than that in the passive viewing tasks described earlier, which might in part reflect the effects of hypermouse concatenation noted above (see also Discussion). Unlike the passive viewing tasks, here we no longer observe a monotonic hierarchical organization of bracket coding strength across visual areas. Nevertheless, significant long-range synchrony among visual regions was still observed, with the majority of dominant interactions (two out of three) consistent with bottom-up information flow (Figure 5e). Also consistent with earlier findings, we observe little to no evidence of bracket coding or its long-range synchrony in HPF.

Our analysis of the IBL data further reproduced the observation that distinct dimensions of sensory information are encoded within different brackets. Using the same temporal dissimilarity analysis, we found significant correlations between activity in higher visual areas, V1, and LGd and the boundaries of real brackets compared with shuffled controls. Temporal multiplexing analysis further revealed significant negative and positive correlations between bracket boundaries and mean decoding accuracy and its temporal derivative, respectively. Consistent with these findings, decoding based on empirically identified brackets outperformed decoding using randomly generated boundaries (Figure 5f).

We also observe weaker, but hierarchically-ordered and partially statistically significant alignments between bracket boundaries and low-frequency LFP oscillations, particularly in the *α*–*β* band in this dataset (Figure 5g). This effect is also specific to the visual hierarchy (not present in HPF), which reinforces its functional relevance. Overall, the consistency of these observations with our findings from the passive viewing tasks in the Allen Institute data supports the generality and robustness of bracket coding as an organizational principle of neuronal population spiking dynamics during visual processing, while also highlighting the need for large datasets in its end-to-end decoding-based discovery.

### Bracket coding in synthetically generated data

To probe the potential generative mechanisms of bracket coding, we next generated synthetic spike trains with parametrically-controlled dynamics and subjected them to the same sweeping–integration pipeline used for real data (Figure 6). Interestingly, we observed the strongest evidence of bracket coding in synthetic data generated by inhomogeneous Poisson processes with dynamic rate profiles that (1) *crossed and changed order* across conditions (trial types) and (2) the times at which these dynamic rates crossed were synchronized across neurons (Figure 6a). This mechanism satisfies both the within-bracket rate coding and the between-bracket temporal coding that signify the bracket coding hypothesis.

**Figure 6.**
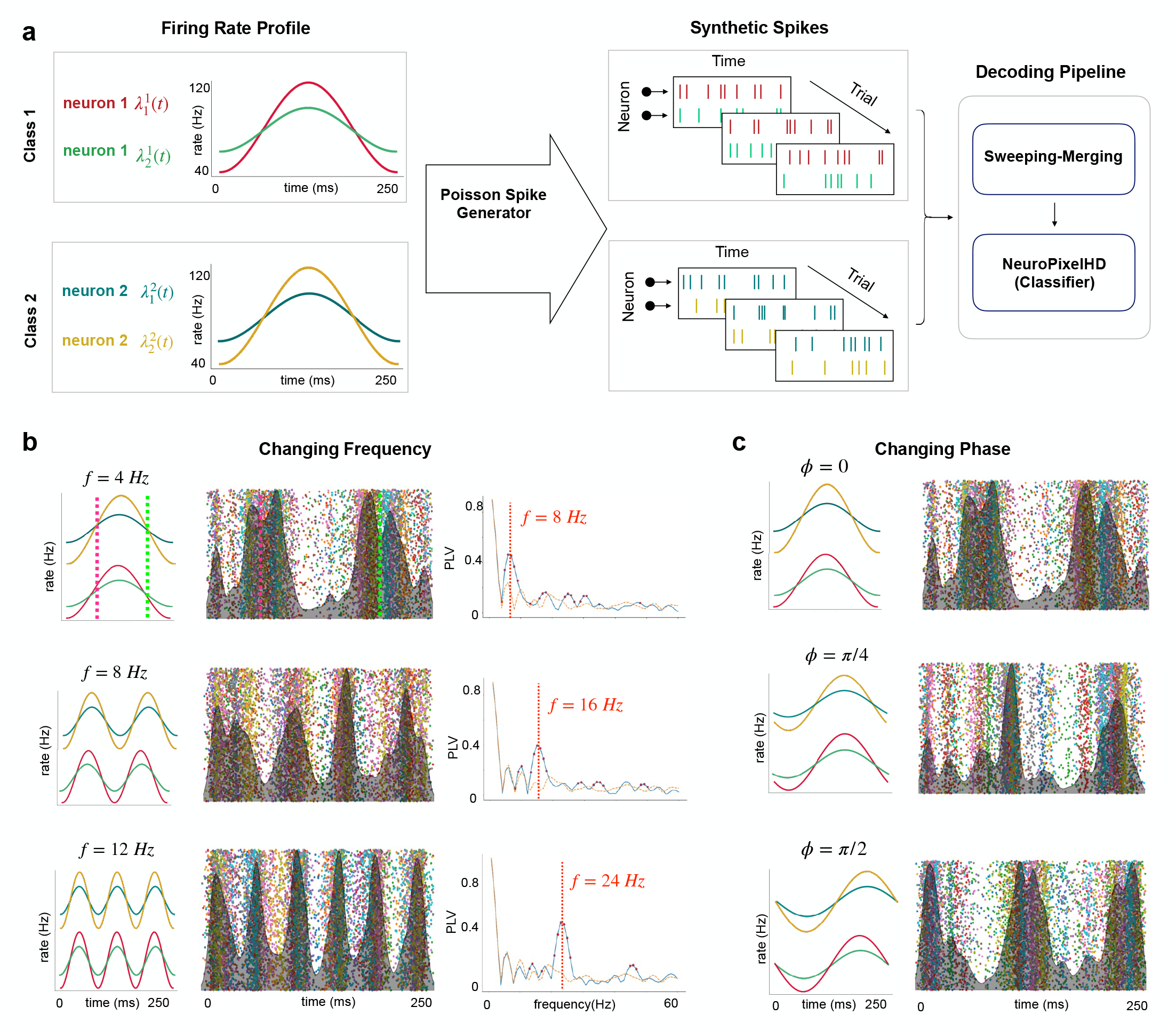
Bracket coding emerges from cross-condition switching and cross-neuron synchrony in synthetic data. (a) Synthetic spike trains were generated using inhomogeneous Poisson processes with dynamic rate profiles that varied across conditions but maintained equal mean rates across neurons. Two neurons and two stimulus conditions were simulated for simplicity. Condition-specific firing-rate trajectories were sinusoidally modulated such that their relative ordering crossed over time, while the timing of these crossings was synchronized across neurons. The resulting spike trains were passed through the same sweeping–integration and decoding pipeline used for real data. **(b)** Effect of modulation frequency. The frequency of the sinusoidal rate modulation was varied while keeping other parameters fixed (sinusoidal rates color-coded as in (a)). Clear bracket boundaries emerged in all cases, while increasing the modulation frequency increased the number of intersections between condition-specific rate trajectories, which in turn increased the number of detected brackets. Vertical lines (4Hz panel) mark the time of switches in the rate profiles for ease of comparison with the trough raster. Details of trough rasters, KDEs of trough distributions, and PLVs (real and shuffled) parallel those in Figures 1-5. Red dots on PLVs show frequencies at which real PLVs are significantly higher than shuffled surrogates (*p <* 0.05, one-sided bootstrap test, FDR corrected for multiple comparisons). **(c)** Effect of modulation phase. Changing the phase of the sinusoidal modulation shifted the timing of the rate intersections and, accordingly, the timing of bracket boundaries.

Using two neurons and two conditions for simplicity, we generated several, parametrically-varying sets of synthetic spiking trajectories satisfying the above two characteristics. Each set consisted of independent sinusoidally modulated Poisson processes that, for each neuron, had the same mean firing rate but different amplitudes of sinusoidal modulation across conditions (see Methods for details). The frequency and phase of the sinusoidal modulation was then parametrically varied across sets, and each set was tested for evidence of bracket coding using the same analysis applied to real data. Across these simulations, bracket boundaries consistently emerged at the intersection points of the dynamic firing-rate trajectories across conditions (Figure 6b). Increasing the frequency of the sinusoidal rate modulation produced a corresponding increase in the number of these intersections, which in turn increased the number of detected brackets with remarkable precision. Further, changing the phase of the sinusoidal modulation proportionately shifted their temporal locations (Figure 6c).

From a mechanistic perspective, these results demonstrate that bracket coding does not require unrealistic discontinuities in spiking statistics or externally imposed boundaries. Instead, brackets can emerge from multivariate dynamic rate profiles as long as the above two properties (cross-condition switching and cross-neuron synchrony) are satisfied. In combination with our analyses of diverse real data, our findings support bracket coding as a novel, potentially optimal, combination of rate-based and temporal-based information encoding in the brain.

## Discussion

### Summary

In this study, we identified bracket coding as a robust mesoscale organizing principle of visual population activity, in which neural responses are partitioned into temporally bounded intervals of relatively stable representation separated by coordinated transitions. Across tasks and datasets, bracket structure was strongest in visual regions across the thalamocortical hierarchy and weak or absent in the hippocampus, indicating its specificity and functional relevance to sensory processing. Cross-correlation analysis of bracket boundaries further revealed a consistent bottom-up temporal ordering, with bracket boundaries in the visual thalamus leading those in V1 which in turn lead those in higher visual areas (HVAs), even as the strength of bracket organization *increased* along the same hierarchy. This pattern suggests that bracket structure is not simply inherited from feedforward sensory input, but is progressively refined and amplified through thalamocortical processing. Consistent with this view, our temporal dissimilarity analysis showed that bracket boundaries are associated with changes in the information carried across neighboring brackets, suggesting that bracket boundaries capture meaningful shifts in representational structure over time. More broadly, this organization is consistent with a temporally structured or multiplexed encoding regime, in which neural populations distribute information across multiple timescales and temporal intervals. The association we observed between bracket coding and low-frequency LFP oscillations further raises the possibility that slow circuit-level rhythms may contribute to the biological generation or coordination of bracket structure, while our simulations point to temporally coordinated switching in dynamic firing-rate profiles across conditions as another potential mechanism contributing to bracket formation.

### Potential mechanisms of bracket formation

While the present study does not resolve the circuit mechanisms underlying bracket coding, our results point to two possible contributors. First, bracket boundaries showed significant phase locking to low-frequency LFP activity, particularly in the theta and alpha–beta bands, indicating that slow mesoscopic network dynamics may help coordinate the timing of bracket transitions. Second, the synthetic-data analyses provide a complementary computational interpretation, where bracket boundaries can emerge consistently at the intersection points of dynamic firing rate trajectories across conditions. Therefore, bracket coding can arise naturally from multivariate dynamic rate codes *provided that* partially synchronized changes in firing-rate structure generate discrete transitions in population decodability. In this view, brackets may reflect the joint action of slow circuit-level coordination and structured rate reconfiguration within neural populations, providing a mechanistic bridge between oscillatory population dynamics and the temporally segmented encoding of sensory information.

### Comparison across datasets

While almost all of our initial findings using the passive view tasks in the Allen Institute dataset replicated during the active discrimination tasks in the IBL dataset, we generally observed weaker effect sizes in the latter. In both cases, bracket coding was specific to visual regions and observed with increasing intensity along the visual pathway, with greatest evidence in higher visual areas (HVAs). Likewise, several core features of bracket organization were reproduced across datasets, including the encoding of distinct dimensions of sensory information across different brackets, and significant cross-region lead–lag relationships that are broadly consistent with a bottom-up thala-mocortical information flow. At the same time, while in the Allen dataset the strength of bracket coding followed a clear hierarchical gradient across visual regions, this hierarchy was less regular and monotonic in the IBL dataset. Several factors likely contribute to this attenuation. First, since the IBL dataset contains fewer trials per condition, we were not able to divide the data into as many independent subtasks as we were able to do for the passive viewing tasks. This reduction in turn limits our ability to reliably estimate the underlying distributions of PLVs in real and shuffled data and thus weakens the statistical separation between the two. Furthermore, unlike the Allen dataset which probed passive sensory responses under relatively controlled conditions, the IBL task involved active perceptual decision-making and motor behavior which introduce additional sources of variability and may partially mask sensory bracket structure. Moreover, the number of neurons available for analysis in IBL was lower after neuron selection, and trial counts were non-uniform across conditions, both of which further reduce statistical power.

Another likely influential feature of the IBL analysis was the construction of hypermice, in which neurons from multiple animals were pooled and treated as a single population. This was done out of computational necessity, since compared with the Allen dataset, the IBL recordings offered a much larger number of animals but fewer neurons and fewer highly informative trials per individual mouse, making single-animal end-to-end decoding substantially less reliable. By aggregating across mice, we increased the effective population size and informational richness available to the decoder. Nevertheless, this procedure is expected to attenuate bracket coding, as only the temporal structures that are shared across all constituent animals can be discovered at the group level. For this reason, the persistence of significant, though weaker, bracket structure in the IBL data is particularly informative and noteworthy. Overall, our comparisons across the two datasets suggest that the data-driven computational procedure needed for the discovery of bracket structure relies on several factors, including conditions of dense sampling, more uniform trial statistics, and passive sensory stimulation.

### Bracket coding beyond stimulus onset response

A strong bracket boundary that we consistently observed across tasks reflects the onset of stimulus-evoked response in the respective thalamocortical areas (Figure 2). While reassuring, this raises the question of whether this marks the *only* significant bracket boundary (followed by purely rate-coded information). To assess this, we repeated PLV analyses after excluding error-curve troughs within the first 70 ms of all trials. As expected, removing the leading bracket boundary contained withing this interval reduced the overall PLV magnitude across visual regions. Nevertheless, we still observed significant evidence of bracket coding in the the remaining trial duration. Across the three passive visual tasks—Gabor patches, natural scenes, and drifting gratings—real PLVs remained significantly higher than shuffled surrogates in visual regions, with particularly strong and significant effects in HVAs and V1 (Supplementary Figure S4). In LGd, the real–shuffled difference was still present but weaker and did not consistently reach significance. This pattern is consistent with the hierarchical organization shown in Figure 4, where bracket-coding strength was lowest in LGd. Thus, after removing the dominant early bracket, the weakest visual region naturally falls below significance, while higher visual cortical areas continue to show robust bracket structure. As in the original analysis, HPF continued to show little to no evidence of bracket coding. Together, these results indicate that the while the leading stimulus-onset bracket boundary contributes substantially to the elevated PLV in real data, it does not fully account for bracket coding which remains evident beyond the onset period.

### PLVs and bracket coding at the subtask level

In the main analysis, we quantified bracket-coding strength by overlaying troughs across independent subtasks and computing the PLV of the aggregated trough distribution. This approach was intended to identify bracket boundaries that are shared across subtasks, and therefore provides a strict measure of global temporal structure across independent subtasks. As a complementary analysis, we also computed the PLV separately within each subtask and then averaged the results, as shown in Supplementary Figure S5. As expected, this subtask-level analysis yielded higher PLV values compared to Figure 4. However, subtask-level PLVs also showed greater and more significant real–shuffled separation than that observed in aggregated troughs, demonstrating the presence of additional subtask-level temporal structure that cannot be explained by rate structure alone. In other words, subtask-level PLVs further show that real population activity contains significant temporal structure beyond that explainable by rate coding, some of which are subtask-specific while others are globally shared across all subtasks.

### Use of frequency-resolved PLV as a measure of temporal alignment

In this study we used frequency-resolved (single-frequency) PLVs as the main measure of temporal alignment between error troughs corresponding to different reference times (RTs) and subtasks (Figure 1d). While such PLVs provide a sensitive measure to capture the distinction between shared temporal structure across RTs and subtasks and broadly-distributed troughs arising from finite-sample variability, they often underestimate the presence of temporal structure that is not necessarily periodic. While the latter can be potentially addressed by using more complex measures, we preferred frequency-resolved PLVs because of their simplicity and interpretability. Further, the sensitivity of frequency-resolved PLVs to periodic structure is also advantageous,as it allows us to quantify the timescales over which bracket coding is strongest. This analysis was further reinforced by the assessment of phase-locking to slow LFP oscillations, where single-frequency phase dynamics were replaced by narrow-band oscillations specific to each subject.

### Absences of bracket coding

While we consistently failed to find evidence of bracket coding in the hippocampal formation (HPF) as a successful demonstration of the specificity of our findings to visual regions, there were other instances were an absence of bracket coding was rather unexpected. One such instance was bracket coding of motor-related dynamics. To further investigate the functional specificity of bracket coding, Using the IBL dataset, we extended our analyses to neuronal populations in both primary and secondary motor areas (MOp and MOs) to decode hypermice’s behavioral choices (left vs. right wheel turn) (Figure S3b). We analyzed the 400ms window preceding movement onset, and constructed two subtasks by independently partitioning left and right movement trials into two non-overlapping subsets of equal size. Nevertheless, in contrast to the encoding of visual information, we did not find significant evidence that motor-related activity is bracket-coded (Figure S3c). A likely contributing factor to this difference is how visual and motor-related information are organized over time. Visual responses are strongly locked to stimulus onset and unfold along a relatively ordered sensory hierarchy, conditions that are favorable for the emergence of temporally aligned transitions and reproducible bracket boundaries. By contrast, movement-related signals are broadly distributed across many regions of the brain and are mixed with choice, engagement, and ongoing behavioral state [42]. Motor population activity is also structured over longer windows of preparatory and movement-related epochs with greater trial to trial variability in timing, even after time-locking to apparent movement onset [10, 15]. Therefore, our observed lack of bracket coding of motor information likely reflects an inherent difference between sensory and motor dynamics where visual coding appears more temporally aligned and hierarchically structured, whereas motor-related information is more distributed and less time-locked to a common external temporal reference.

Another instance were we did not observe evidence of bracket coding was encoding of passive visual information during later intervals of each trial. Using data from the drifting grating task in the Allen Institute dataset, we applied the same sweeping-integration analysis separately to successive 250ms intervals of each trial. Interestingly, unlike the clear evidence of bracket coding that we observed during the first 250ms interval (Figure 2-4), the subsequent intervals did not exhibit reliable bracket structure (Figure S3). This reinforces the idea that the precise temporal structure that lies at the core of bracket coding is primarily present during early-onset processing of visual information, and gradually fades over longer timescales. In other words, our results suggest that spike brackets are most prominent when the neural population is undergoing rapid reorganization of sensory representation time-locked to an external sensory onset, and become weaker over longer timescales where response timing becomes more variable over trials.

### Comparisons with packet coding

Bracket coding bears an important conceptual resemblance to the packet-coding framework proposed by Luczak and colleagues [23], while also differing from it in several key respects. In short, packet coding proposes that cortical responses are organized into transient population events of roughly 50–200ms, within which neurons participate in a broadly conserved sequential pattern and stimulus identity is conveyed by variations in spike timing and firing rate. In contrast, the brackets we identify are shorter, typically tens of milliseconds in duration, and are defined operationally by transitions in decoding structure rather than by sequential spike motifs within the interval itself. Accordingly, our results suggest a coding regime in which representations are relatively stable within brackets and change primarily at their boundaries, whereas in packet coding the fine temporal order within the interval is itself the dominant code. The two views are therefore best seen as complementary rather than competing: packet coding describes a broader, temporally extended motif of cortical population activity, whereas bracket coding identifies a finer-scale segmentation of neural responses into decoding-relevant epochs with comparatively precise boundaries. Furthermore, while bracket coding is currently observed only in stimulus-evoked visual responses, packet coding was proposed as a general principle spanning both spontaneous and sensory-evoked cortical dynamics [23].

### Limitations

This study has a number of limitations. Most importantly, our results are limited to representations of visual information in the first few hundreds of milliseconds after stimulus onset. As noted, we did not observe evidence of bracket coding in later intervals or that of motor-related information, a degree of specificity that needs to be refined in subsequent studies. A further practical limitation of our approach is that the sweeping-integration and decoder-based analyses require relatively large, highquality population datasets with sufficient number of trials, neurons, and stimulus conditions to robustly estimate boundary structure. This may, in turn, limit its use and replication in smaller or noisier datasets. Finally, although our results identify consistent statistical and dynamical signatures of bracket organization, they do not reveal the precise circuit-level mechanisms responsible for generating them. Addressing these limitations will require extending the framework to broader neural systems, richer behavioral paradigms, and experimental designs that can more directly probe mechanisms.

### Conclusions

Our results identify bracket coding as a novel, functionally relevant organization of visual population activity in which neural responses are segmented into bounded intervals of relatively stable representation separated by coordinated transitions. Cross-regional analyses further suggest that bracket formation emerges earliest in sensory thalamus and propagates, while becoming progressively stronger, along the visual hierarchy. Future work will be needed to more precisely map the regions and conditions under which bracket coding arises in neural population activity, and determine the circuit-level mechanisms that give rise to such hybrid encoding structure.

## Methods

### Data

Neural recordings analyzed in this study were obtained from publicly available Neuropixels electrophysiology datasets collected in mice. The primary dataset was the Allen Brain Observatory Visual Coding Neuropixels dataset, which provides large-scale recordings of spiking activity across multiple cortical and subcortical regions during passive visual stimulation [40] In short, awake head-fixed mice viewed a variety of visual stimuli while neuronal activity was simultaneously recorded from hundreds of neurons using high-density Neuropixels probes. We analyzed responses to three stimulus classes: Gabor stimuli, natural scenes, and drifting gratings, which probe different aspects of visual processing. These recordings include activity from several regions along the visual hierarchy, including higher visual areas (HVAs), primary visual cortex (V1), and the visual thalamus (dorsal lateral geniculate complex, LGd). As a control region outside the visual pathway, we also analyzed neurons recorded in the hippocampal formation (HPF). To complement these passive viewing tasks, we also analyzed recordings from the International Brain Laboratory (IBL) Neuropixels dataset, in which mice performed an active visual contrast discrimination task [3]. In short, a visual grating stimulus appeared on either the left or right side of the screen with varying levels of contrast, and mice reported their decision by turning a steering wheel to move the stimulus toward the center of the screen. Data from various brain regions were collected, but only data from HVAs, V1, LGd, and HPF were analyzed in this study. For further details on the respective tasks and data collection procedures, please refer to [40] and [3], respectively.

### Spike preprocessing

Spike times from sorted single units were obtained directly from each dataset. For each trial, spike trains were aligned to stimulus onset and analyzed within a post-stimulus window of 250ms for the Allen Institute and 400ms for the IBL dataset. Spike times were discretized into 1 ms bins, producing a time-resolved spike-count matrix for each trial in which rows correspond to neurons and columns correspond to time bins. Analyses were performed separately for each major brain region, with the corresponding anatomical areas included in each region shown in Table 1. The resulting concatenated population spike trains were provided as input to sweeping integration pipeline described earlier (Figure 1).

**Table 1.** Brain regions analyzed in this study and their constituent areas.

| Region | Constituent Areas |
| --- | --- |
| HVAs | VISal, VISam, VISl, VISpm, VISrl, VISmma, VISli |
| V1 | VISp |
| LGd | LGd |
| HPF | CA1, CA2, CA3, DG, SUB |

### Organization of subtasks

To increase statistical power in distinguishing true brackets from bracket-like non-uniformities that may arise in shuffled trough densities due to finite samples, we constructed multiple ‘subtasks’ under each visual task. Each subtask corresponds to a multi-class classification problem where a subset of stimuli (classes) was selected from all available stimuli within that task and the end-to-end decoder (NeuroPixelHD, see below as well as Figure 1) was trained on trials from that subset only. For the Gabor task, stimuli were originally arranged on a 9 × 9 spatial grid, yielding a total of stimulus classes. We partitioned these into 9 subtasks by uniformly selecting sets of 9 stimuli across the 9 x 9 grid (Figure 2a). For the natural scenes task, stimuli originally consisted of 118 distinct image. These were divided into 10 subtasks where subtask 1 contained images 0, 10, … 110, subtask 2 contained images 1, 11, …, 111, and so on (Figure 2a). For the drifting gratings, stimuli were originally shown at a total of eight drifting directions (0, 45, 90, 135, 180, 225, 270, and 315 degrees) and five temporal drifting frequencies (1, 2, 4, 8, and 15 Hz). We divided these into 10 subtasks by grouping the eight drifting directions into two groups (0-135 and 180-315 degrees) each of which gave rise to five subtasks based on drifting frequency (Figure 2a). Note that in all above cases, subtasks are non-overlapping and cover all available stimuli within that task. For the IBL task this was slightly different, where we first selected the subset of trials with the highest contrast levels (100% and 25%) and other trials were removed from further analysis. This was done to ensure the learning of meaningful patterns by the end-to-end decoder. The remaining trials were partitioned into two subtasks, each consisting of a 100% contrast on one side of the visual field and a 25% contrast on the other (Figure 5a).

### End-to-end decoder (NeuroPixelHD)

The decoding component is implemented using NeuroPixelHD, a hyperdimensional computing (HDC)–based decoder designed in our prior work [36] for the specific purpose of detecting optimal integration windows for NeuroPixel spiking data. The use of HDC makes the internal computations of the decoder (1) averaging free and (2) insensitive to input dimensionality, both of which are critical for unbiased comparisons between pre-decoder integration levels. The structure of NeuroPixelHD consists of two main stages: an encoding stage and a training stage. In the encoding stage, the population spiking data is mapped into a high-dimensional hyperspace, producing hypervectors of a *fixed dimension*. In the training stage, encoded representations of a training subset of trials are used to learn class hypervectors, which are then used to classify encoded representations of test trials via maximum cosine similarity. For additional details on the structure and properties of NeuroPixelHD, see [36].

### Sweeping-integration algorithm

Pre-processed single-unit spike times binned at 1ms resolution were passed through a sweeping-integration algorithm to detect windows of optimal integration. The algorithm operates exclusively along the temporal dimension. For each (fixed) reference time *t*_*r*_ and (moving) limit time *t*_*l*_ within the trial duration, the spike counts of each neuron are integrated (summed) over the window [min(*t*_*l*_, −*t*_*r*_), max(*t*_*l*_, *t*_*r*_)], and the resulting (smaller) array of population spike counts is passed to NeuroPixelHD for decoding. Therefore, as the integration limit moves away from the reference point, spikes within the window of size *w* = |*t*_*r*_ − *t*_*l*_| are cumulatively integrated, while spikes outside the window remain represented at 1ms resolution. The values of *t*_*l*_ and *t*_*r*_ are then varied from 0 to trial length (250ms for Allen and 400ms for IBL data) in increments of 1ms in two nested loops, and the corresponding decoding accuracy was recorded. The decoding accuracies for each *t*_*r*_ (across all *t*_*l*_) are concatenated into a decoding trajectory 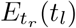, and the process is repeated for each *t*_*r*_. To reduce random fluctuations in the resulting error trajectories, each trajectory is smoothed with a zero-phase 4th-order butterworth lowpass filter (*f*_*c*_ = 30*Hz* in Allen and 20*Hz* in IBL data) and averaged over 50 independent repetitions of the entire procedure with different random seeds. Troughs (local minima) were then detected for each trajectory using and visualized as a raster plot, where each row corresponds to a reference time *t*_*r*_and each dot indicates the location (*t*_*l*_) of a trough.

### Kernel density estimation (KDE) of trough locations

To identify consistent temporal locations at which troughs occur, the trough positions detected for all reference times were pooled and analyzed using a kernel density estimate (KDE) over time. For a given the set of detected trough times 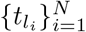, their density was estimated using a Gaussian kernel density estimator, given by

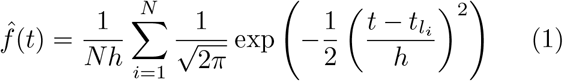

where *h* is the bandwidth parameter controlling the degree of smoothing. The KDE thus provides a continuous representation of trough concentration, the peaks of which define bracket boundaries for each brain region and task.

### Shuffled data

To delineate true temporal structures from apparent structures arising from finite-sample estimates in various analyses, we generated shuffled control datasets whereby spike times were randomly permuted (within each trial) for each neuron, independently across neurons and trials. The same sweeping–integration and subsequent analyses performed on real data were then applied on the resulting shuffled surrogates.

### Computing single-frequency phase-locking values (PLVs)

To quantify the temporal consistency of trough locations across reference times, we computed their PLVs at each frequency *f* ∈ [1, 60] Hz via

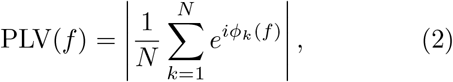

Where

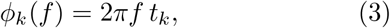

denotes the phase associated with each trough time *t*_*k*_ (measured in seconds), and *N* is the total number of troughs pooled across reference times (and subtasks, when applicable). A larger PLV(*f*) indicates that troughs occur at consistent phases related to a sinusoidal oscillation at *f* Hz, whereas PLV(*f*) ≈ 0 indicates dispersed timing with no consistent alignment.

### Computing PLVs relative to local field potential (LFP)

Similar to single-frequency PLVs described above, alignment of trough locations to the phase of narrow-band LFP oscillations were measured via their relative PLV. For each recording session, LFP signals were obtained for each probe and bandpass filtered into each frequency band using zero-phase finite-impulse-response (FIR) filters. We then segmented and selected LFPs over the duration of trials corresponding to each subtask under consideration and, within each trial, selected LFP channels associated with neurons belonging to the targte brain region. The band-limited LFP signals were averaged across selected channels and trials, resulting in a single representative LFP time series for each region, subtask, and frequency band.

The instantaneous phase of the resulting LFP signal was computed using the Hilbert transform. In short, given the band-pass filtered LFP signal *x*_*f*_ (*t*) we computed the analytic signal

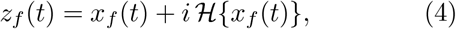

where ℋ{·} denotes the Hilbert transform. The instantaneous phase was then calculated as

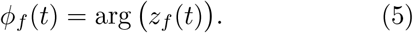

and the corresponding LFP PLV as

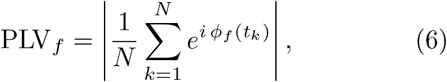

where *N* is the number of trough events pooled across reference times and subtasks.

To assess statistical significance, we generated shuffled controls by randomly permuting trough times. Corresponding LFP PLVs were recomputed for each shuffled realization and combined to obtain a null distribution. LFP PLV values from real data were then compared against this distribution to determine frequency bands and regions exhibiting significant LFP– bracket phase locking.

### Temporal Dissimilarity Analysis

To assess whether bracket boundaries coincide with abrupt changes in decodable information independently of the sweeping–integration procedure, we performed a temporal dissimilarity analysis (cf. Figure 3a), as follows. We defined two adjacent 15 ms sliding windows of fixed duration and moved them across the trial in 1 ms steps. At each time point *t*, we computed a decoding accuracy vector from population spike counts within each window, yielding class-wise accuracy vectors *A*_1_(*t*) and *A*_2_(*t*) for the first and second window, respectively. We then computed the Euclidean distance

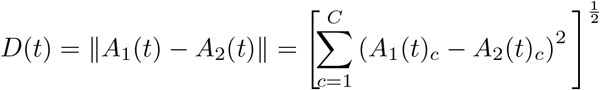

where *C* is the number of classes within that subtask, producing a one-dimensional ‘difference curve’ over time.

To relate changes in decoding structure to bracket timing, we then correlated *D*(*t*) against the kernel density estimate (KDE) of bracket boundary locations obtained from the sweeping–integration procedure, computed separately for each session, brain region, and task. The same analysis was applied to shuffled controls by running the identical pipeline on shuffled spike data.

### Temporal Multiplexing Analysis

To characterize how decoded information evolves over time and how it relates to bracket boundaries, we performed a temporal multiplexing analysis (cf. Figure 3b), as follows. Using a single sliding window (15 ms duration; 1 ms step), we computed class-wise decoding accuracy as a function of time. Let *A*(*t*)_*c*_ denote decoding accuracy for class *c* ∈ {1, …, *C*} at time *t*.

We computed the mean accuracy as

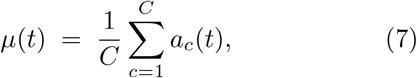

and its temporal derivative as

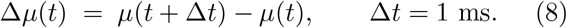

We then linked the dynamics of *μ*(*t*) and Δ*μ*(*t*) to bracket structure by correlating them against the kernel density estimate (KDE) of bracket boundary locations, as described above under temporal dissimilarity analysis.

### Cross-regional lead–lag analysis

To quantify lead–lag relationships in bracket timing across brain regions, we performed a cross-correlation analysis on kernel density estimates (KDEs) of bracket boundaries. Let *g*_1_(*t*) and *g*_2_(*t*) denote the KDEs of two regions, with their dominant peaks marking respective bracket boundaries. We computed their crosscorrelation function

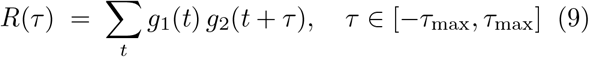

where *τ*_max_ = 30 ms for both Allen and IBL data. The optimal lag estimate for this region pair was then computed as

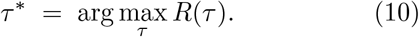

Note that under this convention, *τ*^*\**^ *>* 0 indicates that *g*_1_(*t*) is shifted *later* relative to *g*_2_(*t*) (i.e., boundaries in region 1 tend to lag those in region 2), whereas *τ*^*\**^ *<* 0 indicates the opposite ordering.

This procedure was repeated across subjects and subtasks, yielding a distribution of *τ*^*\**^ values for each region pair. To test for a consistent directional relationship, we assessed whether the median *τ*^*\**^ differed from zero using a Wilcoxon signed-rank test, and applied the same analysis to shuffled controls to establish a baseline in the absence of structured cross-regional timing.

### Selection of mice and hypermice in IBL data

A total of 114 unique mice were included in analyses based on IBL data. Region-specific subsets comprised 51 mice from higher visual areas (55 recording sessions), 32 from V1 (39 recording sessions), 20 from thalamic regions (24 recording sessions), 84 from hippocampal regions (112 recording sessions), 19 from primary motor area (25 recording sessions) and 25 from secondary motor area (31 recording sessions). Additional sessions reflect repeated recordings from the same animals. Sessions were included if each class contained more than 15 trials and fewer then five ‘empty’ trials (i.e., trials with no observable activity). Analyses were restricted to well-isolated neurons, selected based on standard quality-control criteria (see [3] for details). Empty trials primarily arose in recordings comprising such neurons.

A central component of our analysis is the construction of hypermice, which serve as the primary framework for capturing population-level neural structure across multiple mice and recording sessions. To construct a hypermouse, we aggregated neuronal activity from random groups of 15 mice and sessions, forming a unified population-level representation. The choice of 15 sessions per hypermouse reflects a balance between representational coverage and statistical robustness (Figure 5c). For each brain region and subtask, groups of 15 mice were randomly generated and neuronal activity was pooled (concatenated) among them across the neuron dimension. To allow for such pooling, we inevitably reduced (subsampled) the number of trials available for each mouse to the minimum number of trials available among the group. The concatenated data was then used in all subsequent analyses as described above.

### Synthetic data generation

We constructed synthetic spike trains consisting of two neurons and two stimulus conditions (classes), as follows. For each neuron and condition, spiking activity was generated as an inhomogeneous Poisson process whose instantaneous firing rate followed a sinusoidally modulated profile over the 250-ms trial window. All neurons shared the same baseline firing rate, while the amplitude of the sinusoidal modulation varied across neurons and conditions. To introduce complementary information across neurons, the amplitudes were arranged such that the modulation amplitudes were swapped across conditions (classes). In other words, if neuron 1 exhibited a larger modulation amplitude in condition 0, the same amplitude was assigned to neuron 2 in condition 1, and vice versa. This construction ensured that the dynamic firing-rate trajectories across conditions crossed during the trial while remaining synchronized across neurons.

Formally, the firing-rate function for neuron *i* under condition *c* was defined as

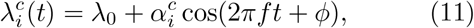

where *λ*_0_ is the baseline firing rate, 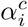 is the modulation amplitude, *f* is the modulation frequency in Hz, and *ϕ* is the phase offset. For each parameter configuration, spike trains were generated by sampling from an inhomogeneous Poisson process with rate 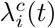 and were analyzed using the same sweeping– integration pipeline applied to real neural recordings.

### Statistical analyses

All statistical analyses were performed using the `scipy` and `statsmodels` libraries in Python. For comparisons between real and shuffled PLVs, a one-sided Wilcoxon signed-rank test (*H*_1_: real *>* shuffled) was employed. Resulting p-values were corrected for multiple comparisons across frequencies using the Benjamini–Hochberg procedure to control the false discovery rate (FDR). For temporal dissimilarity and multiplexing analyses, Pearson correlation coefficients (*r*) were computed between filtered spiking activity and the kernel density estimate (KDE) of bracket boundaries for both real and shuffled data. For temporal dissimilarity, the resulting correlation coefficients were compared using a onesided Wilcoxon signed-rank test (*H*_1_: real *>* shuffled), whereas for multiplexing analysis, a two-sided Wilcoxon signed-rank test was used. Temporal coordination between bracket boundaries and LFP oscillations was evaluated using a one-sided Wilcoxon signed-rank test to determine whether observed PLVs exceeded those from shuffled surrogates. Significance of cross-correlations across regions were also assessed using the Wilcoxon signed-rank test. All statistical significances were evaluated at *α* = 0.05, with significance levels denoted as \**p <* 0.05, * * *p <* 0.01, and * * \**p <* 0.001.

### Computational resources

All analyses were performed on the High-Performance Computing Center (HPCC) cluster at the University of California, Riverside. Data processing, simulations, and decoding analyses were implemented in Python and executed on both CPU and GPU compute nodes using the Slurm workload manager for job scheduling and parallel execution. Depending on the dataset and analysis configuration—including task type, number of trials, and trial duration (250 ms for Allen Neuropixels recordings and 400 ms for IBL datasets)— individual jobs typically used 2 CPU cores and required between 4 and 32 GB of memory.

### Graphic illustrations

Schematic task illustrations in Figures 1a and 5a,b were partially created using BioRender.com.

## Code availability

The code used for the analyses presented in this study is publicly available at https://github.com/nozarilab/2026Samiei_Bracket_Coding.

## Funding

This work was supported in part by the NSF CAREER Award 2239654, the NIH NINDS Award R01-NS-148491, and the DARPA INSPIRE Award HR0011-25-3-0153.

## Competing interests

The authors declare no competing interests.

## Supplementary Figures

**Supplementary Figure S1.**
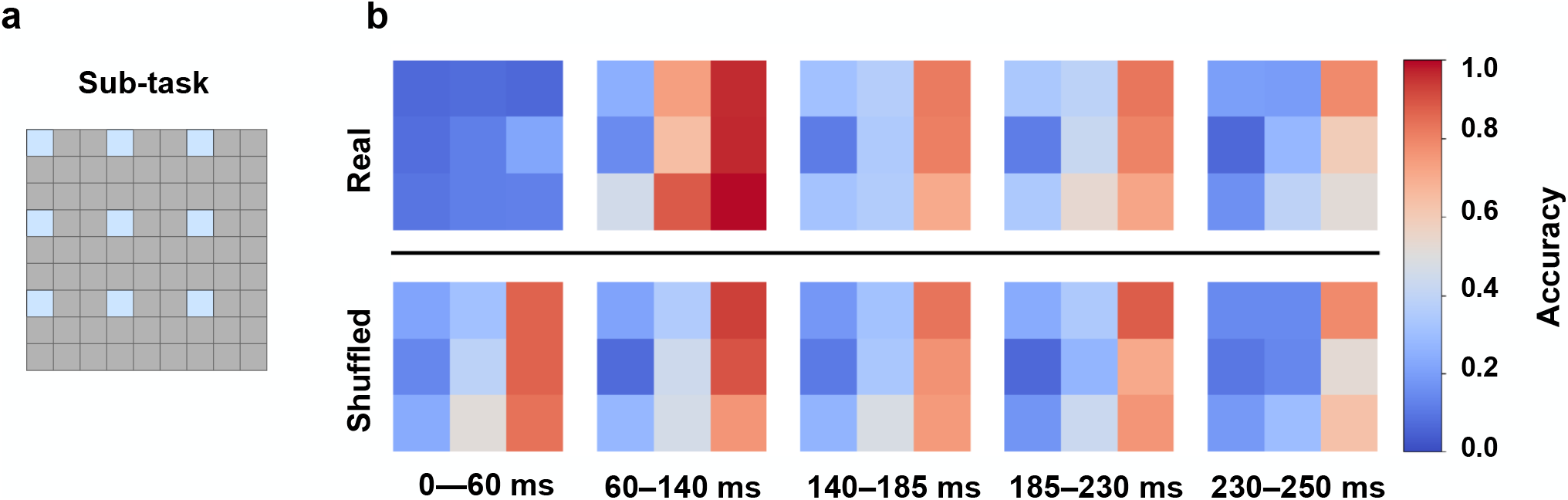
Class-specific decoding profiles across extracted brackets in a representative Gabor subtask. **(a)** Schematic of the representative subtask used for this analysis. **(b)** Heat maps showing the decoding accuracy profiles of these selected classes across brackets extracted from the real population activity. Each heat map summarizes class-wise decoding performance when the decoder was provided only with spike information integrated within a single bracket. The color in each heat map indicates the fraction of correctly classified trials for each class within that bracket. In the real data, the decoding profile of individual classes changes substantially from one bracket to another, indicating that the decodability of a given class evolves over time. In the shuffled data, although some classes remain easier to decode than others, the temporal changes in class-wise decoding profiles are more uniform.

**Supplementary Figure S2.**
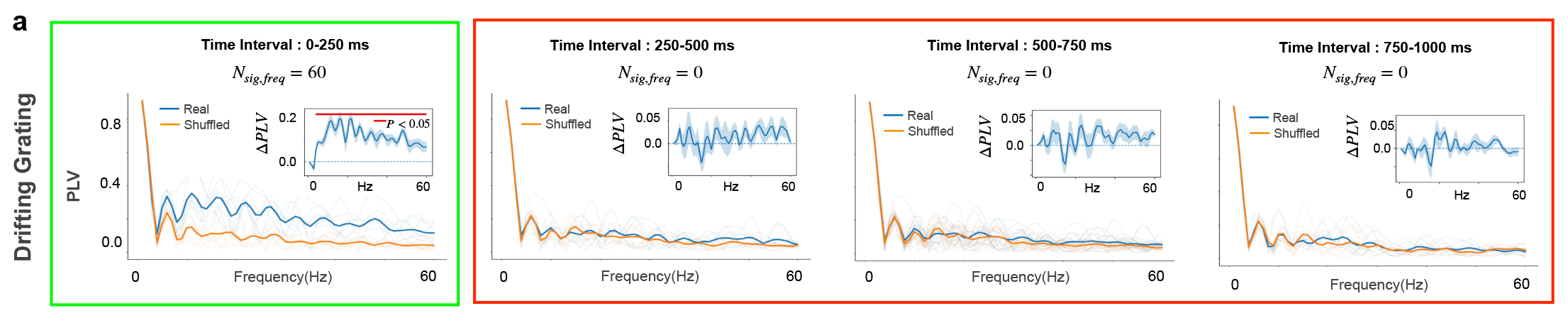
Bracket coding is restricted to early stimulus-evoked response in drifting gratings. **(a)** Sweeping–integration analysis of drifting-grating responses performed separately in consecutive 250ms windows across the first second of stimulus presentation. For each interval, phase-locking values (PLVs) between trough times and sinusoids of different frequencies are shown for the real data (blue) and temporally shuffled surrogates (orange), together with the corresponding ΔPLV spectra (insets). Clear bracket structure was observed only in the first 0–250ms interval, which showed a larger PLV in the real data than in the shuffled data and a nonzero number of significant frequencies (*N*_sig,freq_ = 60), whereas the subsequent intervals (250–500, 500–750, and 750–1000 ms) showed no significant frequencies (*N*_sig,freq_ = 0).

**Supplementary Figure S3.**
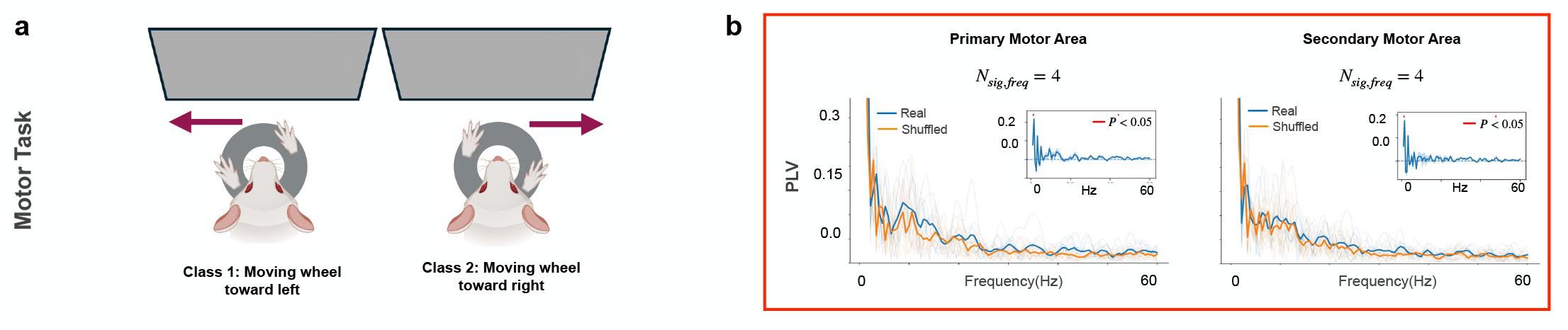
Bracket coding of movement is present but weak in motor regions. **(a)** Schematic of motor choice decoding in the IBL dataset. Neural activity from motor regions was used to decode the animal’s upcoming behavioral choice, defined as wheel movement toward the left or toward the right. The analysis was performed on the 400 ms period immediately preceding movement onset. Two subtasks were defined by independently splitting left and right movement trials into two equal halves. **(b)** PLV spectra for neuronal populations in primary motor area (MOp) and secondary motor area (MOs), shown for real data (blue) and temporally shuffled surrogates (orange), with corresponding ΔPLV spectra in the insets. In contrast to visual regions, motor populations exhibited weak bracket structure, with only a small number of significant frequencies in both MOp and MOs (*N*_sig,freq_ = 4). These results suggest that bracket coding is not a general feature of all task-engaged populations, but is strongest in sensory responses that are tightly aligned to stimulus onset and is substantially weaker in motor-related activity associated with action execution.

**Supplementary Figure S4.**
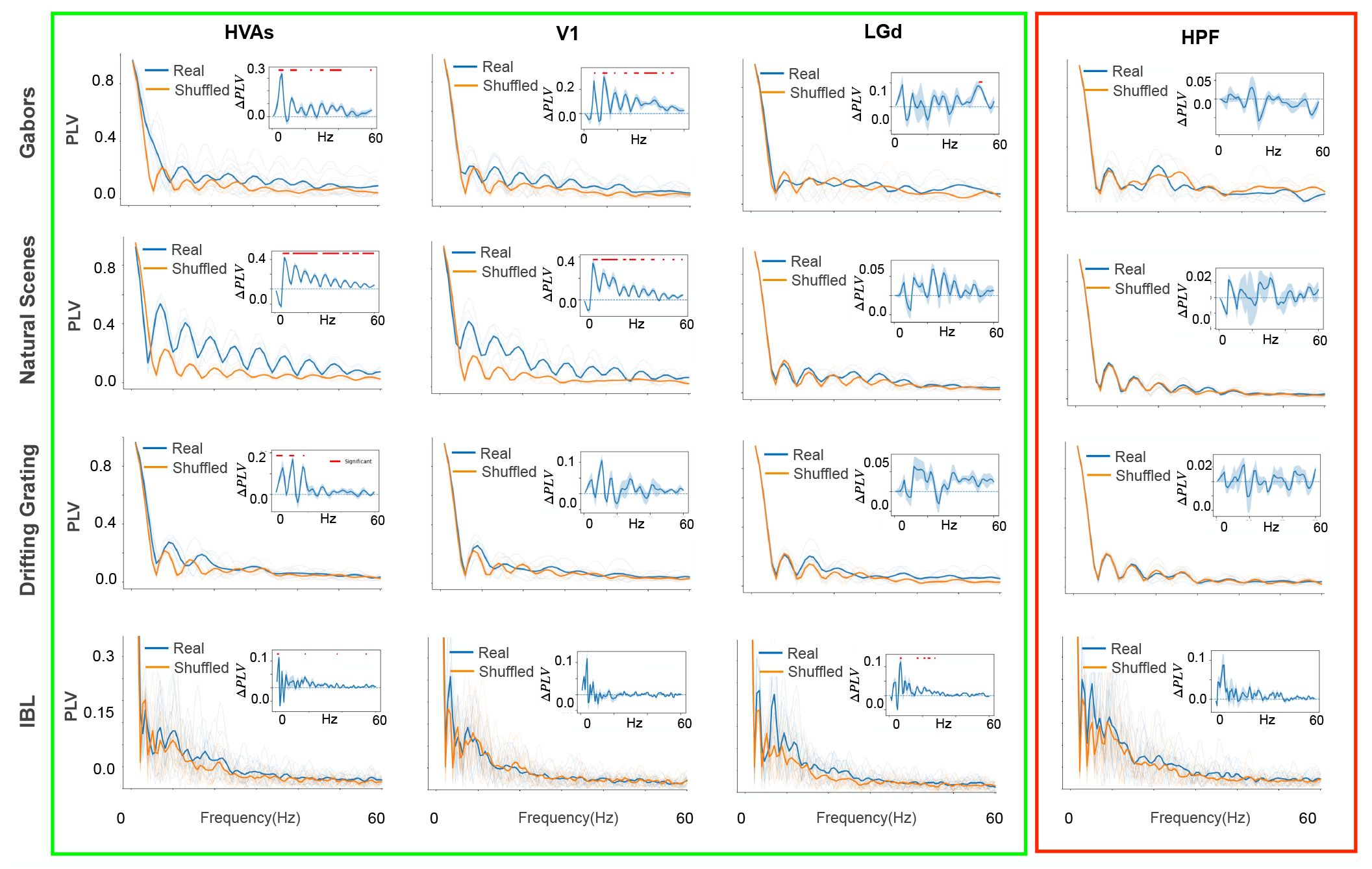
Persistence of bracket coding beyond the stimulus-onset response. To assess the contribution of the first bracket boundary, marking the arrival of stimulus-evoked response in each region at about 50-70ms post stimulus onset, we repeated the PLV analysis after excluding troughs occurring within the first 70ms of each trial duration. Resulting PLV spectra are shown for real data (blue) and temporally shuffled controls (orange) across tasks and regions. Details parallel those in Figure 2 in the main text. As expected, removing the dominant early bracket reduced the overall PLV magnitude across visual regions. However, real PLVs remained higher than shuffled PLVs, particularly in HVAs and V1, indicating that bracket structure persists beyond the earliest stimulus-onset response. LGd showed a weaker real–shuffle separation that did not consistently reach significance, which is consistent with its lower bracket-coding strength in the visual hierarchy (Figure 4a).

**Supplementary Figure S5.**
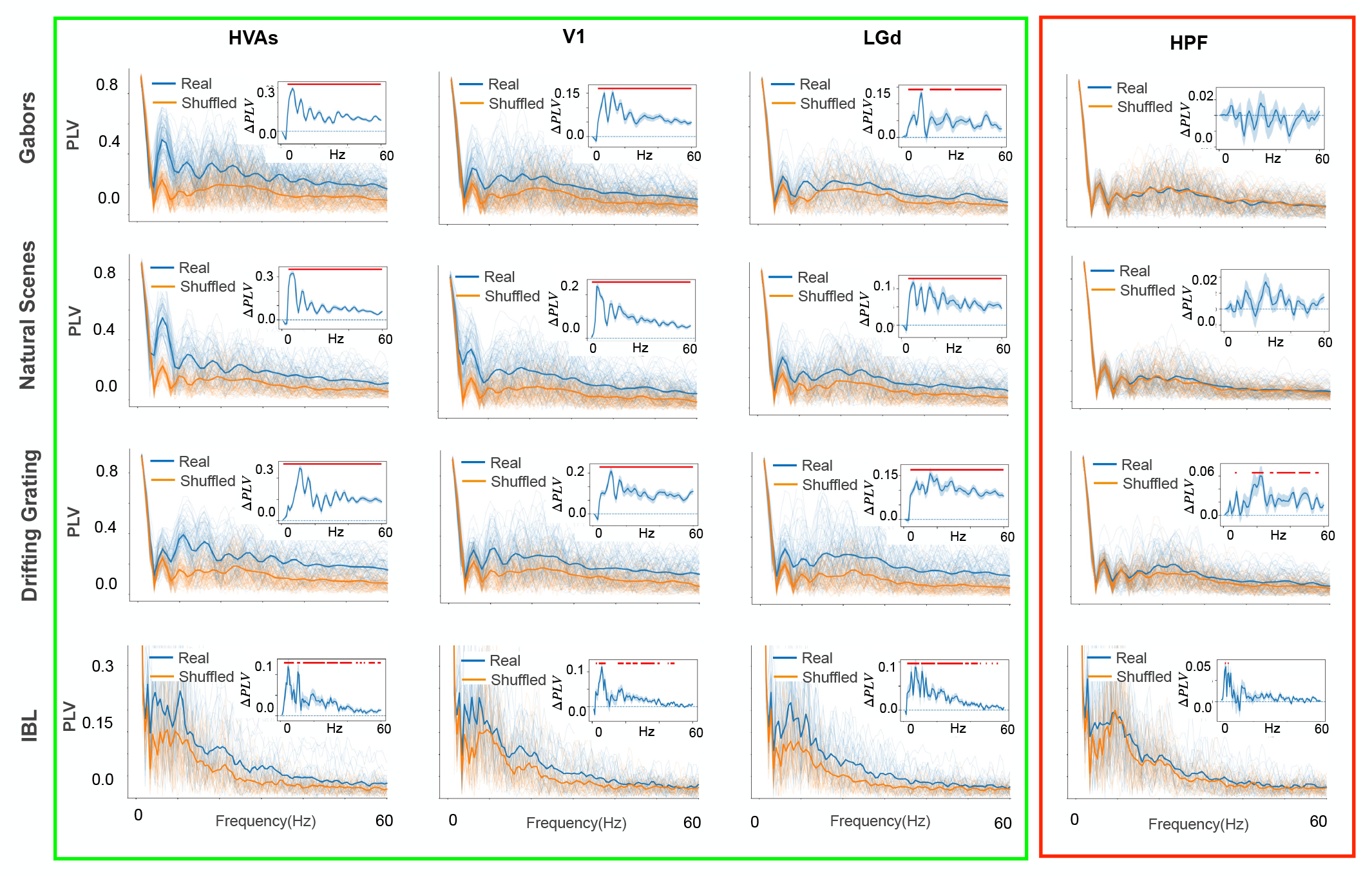
Subtask-specific temporal structure. Complementary to the aggregated-trough PLV analyses in the main text (cf. trough rasters in Figure 2), we also computed PLVs for each subtask and then averaged the results, shown here for real data (blue) and temporally shuffled controls (orange). Details parallel those in Figure 2. As expected, mean subtask-level PLVs are generally higher than aggregated-trough counterparts. However, notably, the former also showed—almost exclusively in visual regions—stronger and more significant separation between real PLVs and shuffled surrogates, compared to the same separation in trough-aggregated counterparts. This result supports the interpretation that the elevated subtask-level PLVs in real data reflects temporal structure beyond what can be explained by firing rates alone.

## Notes

### Competing Interest Statement

The authors have declared no competing interest.

### Summary of Updates

This version of the manuscript has been revised to update the 'title' of the manuscript.

